# Non-invasive surveys of mammalian viruses using environmental DNA

**DOI:** 10.1101/2020.03.26.009993

**Authors:** Niccolò Alfano, Anisha Dayaram, Jan Axtner, Kyriakos Tsangaras, Marie-Louise Kampmann, Azlan Mohamed, Seth T. Wong, M. Thomas P. Gilbert, Andreas Wilting, Alex D. Greenwood

**Affiliations:** Department of Ecological Dynamics, Leibniz Institute for Zoo and Wildlife Research, Alfred-Kowalke-Str. 17, 10315 Berlin, Germany; Department of Biodiversity & Molecular Ecology, Fondazione Edmund Mach, Research and Innovation Centre, Via Edmund Mach 1, 38010 San Michele All’adige (TN), Italy; Department of Wildlife Diseases, Leibniz Institute for Zoo and Wildlife Research, Alfred-Kowalke-Str. 17, 10315 Berlin, Germany; Charité-Universitätsmedizin Berlin, corporate member of Freie Universitäts Berlin and Humboldt-Universität of Berlin, Institut für Neurophysiologie, Berlin, Germany; Department of Translational Genetics, The Cyprus Institute of Neurology and Genetics, Iroon Avenue 6, Agios Dometios 2371, Cyprus; Section of Forensic Genetics, Department of Forensic Medicine, Faculty of Health and Medical Sciences, University of Copenhagen, Denmark; WWF-Malaysia, PJCC, 46150 Petaling Jaya, Selangor, Malaysia; The GLOBE Institute, University of Copenhagen, Øster Farimagsgade 5A, 1352 Copenhagen, Denmark; University Museum. NTNU, 7491 Trondheim, Norway; Department of Veterinary Medicine, Freie Universität Berlin, Robert-von-Ostertag-Str. 7-13, Berlin 14163, Germany

**Keywords:** environmental DNA (eDNA), hybridization capture, leeches, non-invasive samples, viral diversity

## Abstract

1. Environmental DNA (eDNA) and invertebrate-derived DNA (iDNA) have been used to survey biodiversity non-invasively to mitigate difficulties of obtaining wildlife samples, particularly in remote areas or for rare species. Recently, eDNA/iDNA have been applied to monitor known wildlife pathogens, however, most wildlife pathogens are unknown and often evolutionarily divergent.
2. To detect and identify known and novel mammalian viruses from eDNA/iDNA sources, we used a curated set of RNA oligonucleotides as viral baits in a hybridization capture system coupled with high throughput sequencing.
3. We detected multiple known and novel mammalian RNA and DNA viruses from multiple viral families from both waterhole eDNA and leech derived iDNA. Congruence was found between detected hosts and viruses identified in leeches and waterholes.
4. Our results demonstrate that eDNA/iDNA samples represent an effective non-invasive resource for studying wildlife viral diversity and for detecting novel potentially zoonotic viruses prior to their emergence.

## INTRODUCTION

Emerging infectious viruses increasingly threaten human, domestic animal and wildlife health (Johnson et al. 2019). Sixty percent of emerging infectious diseases in humans are of zoonotic origin (Jones et al. 2008). Wildlife trade and consumption of bushmeat, especially in Africa and Asia, have played a role in pathogen spillovers into human populations (Pruvot et al. 2019; Swift et al. 2007). Wildlife markets may have facilitated the spillover of pandemic SARS-CoV-2 to humans (Zhou et al. 2020). The 2002–2003 SARS-CoV outbreak (Drosten et al. 2003), the Ebola outbreak in West Africa (Leroy et al. 2009) and the global emergence of HIV (Sharp & Hahn 2011) have all been linked to wildlife trade and bushmeat consumption. Early detection of novel infectious agents in wildlife is key to emergence prevention. However, identification, surveillance and monitoring of emerging viruses using direct sampling of wildlife often requires enormous sampling investment, particularly for viruses that have low prevalence (Hoye et al. 2010). For example, 25,000 wild birds were sampled in Germany to detect avian influenza prevalence below 1% (Wilking et al. 2009). Similarly, sampling of over 8,157 animals in Poland was required to determine an 0.12% prevalence of African swine fever virus (ASF) (Śmietanka et al. 2016). Developing comprehensive viral surveillance and discovery methods remains challenging and is often hindered by access to free ranging wildlife.

Using non-invasive nucleic acid sources, such as environmental DNA (eDNA) and invertebrate-derived DNA (iDNA), can mitigate difficulties of obtaining wildlife samples under remote field conditions or in developing countries to complement invasive sampling or replace it when invasive sampling is not possible. So far eDNA and iDNA have been mainly used to survey biodiversity, for instance water eDNA has been used to survey mammalian biodiversity at African waterholes (Seeber et al. 2019) and terrestrial haematophagous leeches (iDNA) have been used to survey mammals in Southeast Asian tropical rainforests (Abrams et al. 2019; Schnell et al. 2018; Tilker et al. 2020). Water is ubiquitous in most ecosystems, and, among invertebrates, leeches are abundant and can be easily collected in many tropical rainforests, many of which are major hotspots of emerging infectious diseases (Daszak et al. 2000). Recently, eDNA/iDNA sources have also been applied to monitoring specific wildlife pathogens (Gogarten et al. 2019; Mosher et al. 2017). However, previous studies have focused on known pathogens, whereas most wildlife pathogens are uncharacterized and unknown, particular in tropical areas where emerging viruses are particularly frequent (Jones et al. 2008). To identify and characterize novel mammalian viruses from eDNA (waterhole water and sediment from Tanzania and Mongolia) and iDNA (haematophagous terrestrial leeches from Malaysia) sources, we designed a hybridization capture system using a curated set of RNA oligonucleotides based on the ViroChip microarray assay (Chen et al. 2011) as baits to target every known mammalian viral genome (Fig. 1). Multiple known and novel viruses were identified from both eDNA and iDNA samples including a novel divergent coronavirus strain demonstrating that eDNA/iDNA samples can be used to survey known and unknown viral diversity.

**Figure 1:**
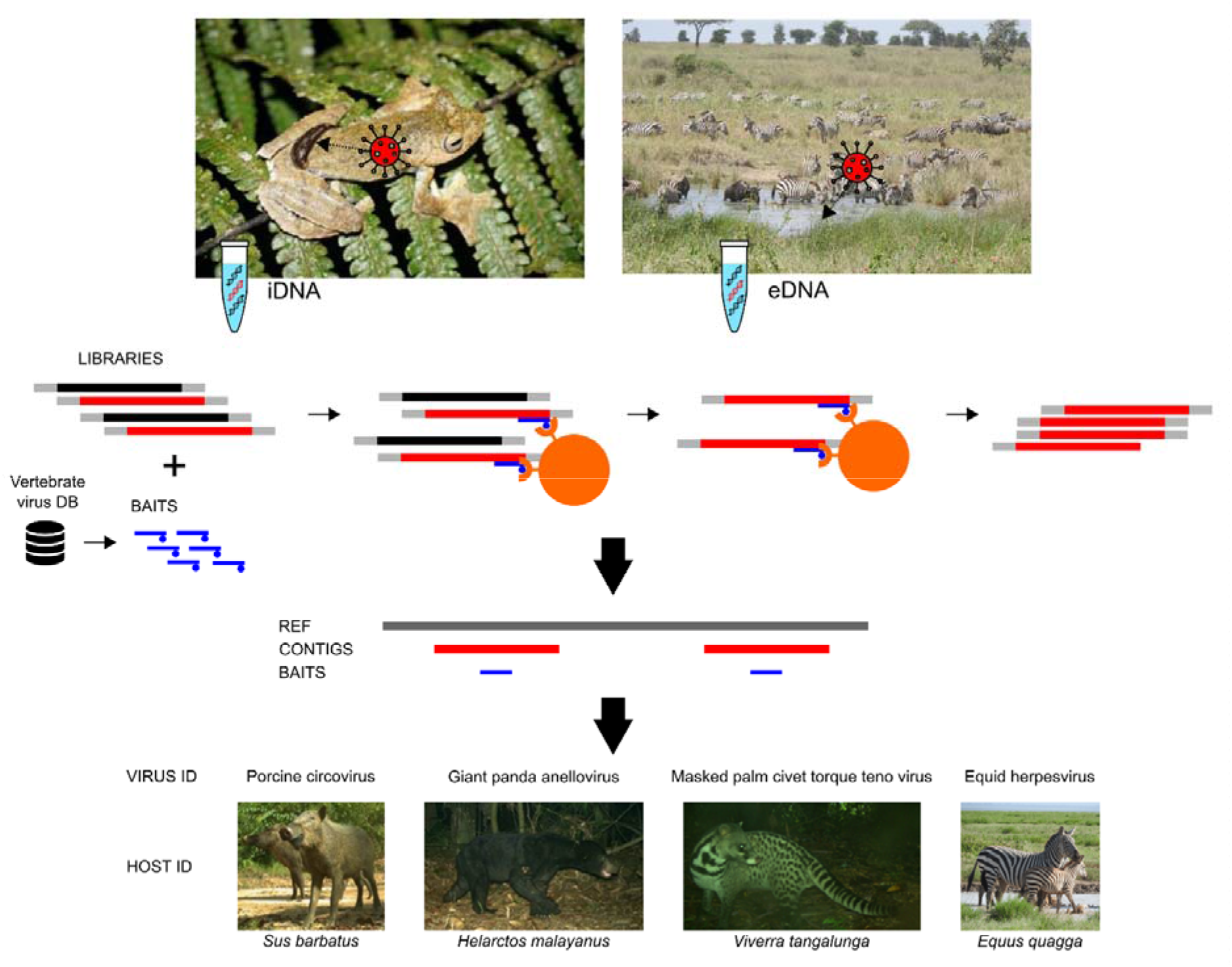
Viral screening of vertebrate viruses from leech iDNA and waterholes eDNA using hybridization capture. In the upper left, a leech feeding on a frog in a rainforest of Vietnam (courtesy Andrew Tilker; Leibniz-IZW). In the upper right, an African waterhole in Tanzania (courtesy Peter Seeber; Leibniz-IZW). In the middle, the hybridization capture protocol: libraries produced from RNA and DNA of leech bloodmeals or waterhole surface water and sediments are hybridized to biotinylated viral RNA baits and non-target material is washed away. The captured molecules are sequenced, reads assembled into contigs and mapped to reference viral genomes. Viral identity is paired with host identity determined either by mammalian metabarcoding of the leech samples, or by observation of waterhole usage.

## MATERIAL AND METHODS

### Sample collection

#### Leeches

Two types of terrestrial leeches, tiger leeches (*Haemadipsa picta*) and brown leeches (*Haemadipsa zeylanica*) were collected from February to May 2015 in the Deramakot Forest Reserve in Sabah, Malaysian Borneo as described in Abrams et al. (2019) and Axtner et al. (2019). All leeches of the same type (tiger or brown) from the same site and occasion were pooled and processed as one sample. Number of leeches ranged from 1 to 77 per pool (median= 7). Samples were stored in RNA fixating saturated ammonium sulfate solution and exported under the permit ‘JKM/MBS. 1000-2/3 JLD.2 (8)’ issued by the Sabah Biodiversity Council. A total of 68 pools (L1-L68) were selected for viral capture to maximize representation of host wildlife species identified from bloodmeals (Axtner et al. 2019).

#### Sediment and water

In February, June, July and October 2016 samples were collected from the Serengeti National Park (ca. 2.2° S, 34.8° E) Tanzania from waterholes. In October 2015 samples were collected from South East Gobi (45.5905°N, 107.1596 °E) and in between June – July 2016 samples were collected from Gobi B (45.1882°N, 93.4288°E) in Mongolia. At each waterhole, 50 ml of water was passed through a 0.22 μm Sterivex filter unit (Millipore) using a disposable 50-ml syringe to remove debris from water. In addition, 25 g of the top 1-3 cm of sediment was collected at each waterhole. The samples were stored on ice packs during the respective field trip, and frozen at −20 °C. In total water filtrate and sediment samples were sampled at 12 waterholes, six respectively from Mongolia and Tanzania. For each sample, 32 ml of water filtrate was ultra-centrifuged at 28,000 rpm for 2 hours to pellet DNA and viral particles. The supernatant was then removed, the pellet re-suspended in 1 ml of cold phosphate-buffered saline (PBS) (pH 7.2) (Sigma-Aldrich) and left at 4 °C overnight.

### Preparation of samples and nucleic acid extraction

#### Leeches

Leeches were cut into small pieces with a new scalpel blade and lysed overnight (≥12 hours) at 55°C in proteinase K and ATL buffer at a ratio of 1:10; 0.2 ml per leech. Nucleic acids were extracted from leech samples using the DNeasy 96 Blood & Tissue kit (Qiagen) (see Axtner et al. 2019 for further details).

#### Water and sediment

500 μl of the centrifuged filtrate was used to extract viral nucleic acids using the RTP^®^ DNA/RNA Virus Mini Kit (Stratec biomedical) with the following modifications: 400 μl of lysis buffer, 400 μl of binding buffer and 20 μl of proteinase K and carrier RNA were used per sample. Samples were eluted in 60 μl. The NucleoSpin Soil kit (Macherey-Nagel) was used to extract DNA/RNA from sediment. 500 mg of soil was extracted according to the manufactures protocol using an elution volume of 100 μl.

### Positive control

As a positive control medical leeches (*Hirudo* spp.) were fed human blood spiked with four viruses (Kampmann et al. 2017). Two RNA viruses, influenza A and measles morbillivirus, and two DNA viruses, bovine herpesvirus and human adenovirus were used (see Kampmann 2017) for details).

### Library Preparation

The RNA was reverse transcribed using SuperScript III and IV (Thermo Fisher Scientific) with random hexamers prior to second-strand synthesis with Klenow fragment (New England Biolabs). The resulting double-stranded cDNA/DNA mix was sheared to an average fragment size of 200 bp using a M220 focused ultrasonicator (Covaris). Sheared product was purified using the ZR-96 DNA Clean & Concentrator-5 kit (Zymo). Dual-indexed Illumina sequencing libraries were constructed as described by Meyer and Kircher (2010) with the modifications described in Alfano et al. (2015). Each library was amplified in three replicate reactions to minimize amplification bias in individual PCRs. The three replicate PCR products for each sample were pooled and purified using the MinElute PCR Purification Kit (Qiagen). Negative control libraries were also prepared from different stages of the experimental process (extraction, reverse transcription, library preparation and index PCR) and indexed separately to monitor any contamination introduced during the experiment. Amplified libraries were quantified using the 2200 TapeStation (Agilent Technologies) on D1000 ScreenTapes.

### RNA oligonucleotide Bait Design

The targeted sequence capture panel was designed based on the oligonucleotide probes represented on the Virochip microarray (Wang et al. 2002). The Virochip is a pan-viral DNA microarray comprising the most highly conserved 70-mer sequences from every fully sequenced reference viral genome present in GenBank, which was developed for the rapid identification and characterization of novel viruses and emerging infectious disease. We retrieved the viral oligonucleotides from the 5th generation Virochip (Viro5) (Yozwiak et al. 2012), which are publicly available at NCBI’s Gene Expression Omnibus (GEO) repository, (https://www.ncbi.nlm.nih.gov/geo/query/acc.cgi?acc=GPL13323). This platform includes ~17,500 oligonucleotides (70-mer nucleotides) derived from ~2,000 viral species. We excluded sequences from bacteriophage, plant viruses, viral families infecting only invertebrates and endogenous retroviruses. We included viruses that could have both vertebrate and invertebrate hosts, such as vertebrate viruses with insect vectors. Exogenous retroviruses were represented but murine leukemia viruses were removed since they can cross enrich endogenous retroviruses which can represent large portions of several vertebrate genomes and mask rarer viral sequences. Control oligonucleotides included in the Virochip, such as those from human genes, yeast intergenic sequences, and human papilloma virus sequences present in HeLa cells were also removed. Ninety-two 70-mer oligonucleotides covering (spaced end-to-end) the entire *pol* and *gB* genes of Equine herpesvirus 1 (EHV-1) were included as PCR screening of the water samples indicated they were positive for this virus (data not shown). The resulting 13,532 oligonucleotides were examined for repetitive elements, short repeats, and low complexity regions, which are problematic for probe design and capture, using RepeatMasker. Repetitive motifs were identified in 234 oligonucleotides, which were removed. The final targeted sequence capture panel consisted of 13,298 unique sequences which were synthesized (as a panel of biotinylated RNAs) at MYcroarray.

### Viral Enrichment Strategy and Sequencing

In-solution target enrichment via hybridization-based capture was performed according to the manufacturer’s protocol (MYbaits^®^ custom targeted enrichment, MYcroarray), with the following modifications for likely partially degraded samples with an expected low target viral content: 50uL Dynabeads^®^ M-270 Streptavidin beads (Invitrogen) instead of 30 uL Dynabeads^®^ MyOne^™^ Streptavidin C1 (Invitrogen); hybridization, bead-bait binding, and wash steps temperature set to 60°C; 48 hours hybridization time; 200 ng baits per reaction; 10 μL indexed library inputs. For capture, libraries generated from pooled leeches consisting of more than 16 individuals were captured individually, while libraries generated from pools of fewer individuals were combined to have a comparable number (15-20) of leeches per capture. This was done in order to ensure that each individual leech represented in each library was allocated enough bait for capture. Water and sediment samples were pooled in groups of two, with sediment and water pooled separately. Per pooled sample, 500 ng of baits were used to ensure enough bait for each sample. The enriched libraries were re-amplified as described in Alfano et al. (2016). The re-amplified enriched libraries were purified using the MinElute PCR Purification Kit (Qiagen), quantified using the 2200 TapeStation (Agilent Technologies) on D1000 ScreenTapes and finally pooled in equimolar amounts for single-read sequencing on two lanes of an Illumina NextSeq 500 with the TG NextSeq^®^ 500/550 High Output Kit v2 (300 cycles).

### Data analysis and bioinformatics pipelines

A total of 219,580,903 sequence reads 300 bp long were generated (average: 3,181,781 single reads per sample; standard deviation [SD]: 1,481,098) (Suppl. Tab. 1) and sorted by their dual index sequences. Cutadapt v1.16 and Trimmomatic v0.36 were used to remove adapter sequences and low-quality reads using a quality cutoff of 20 and a minimal read length of 30 nt. After trimming, 97% of the sequences were retained. Leech reads were removed from the dataset by alignment to the *Helobdella robusta* genome v1.0 (assembly GCA_000326865.1), which is the only complete genome of Hirudinea available in GenBank, and all leech sequences from GenBank (4,957 sequences resulting from “Hirudinea” search) using Bowtie2 v2.3.5.1 (Langmead & Salzberg 2012). This left 81% of the original reads (Suppl. Tab. 1). Then, rRNA reads were removed using SortMeRNA (Kopylova et al. 2012), leaving 75% of the original reads (Suppl. Tab. 1). The filtered reads were *de novo* assembled using both Spades v3.11.1 (Bankevich et al. 2012) and Trinity v2.6.6 (Grabherr et al. 2011) assemblers. The obtained contigs were pooled and clustered to remove duplicated or highly similar sequences using USEARCH v11.0.667 (R. C. Edgar 2010) with a 90% threshold identity value. The centroids were then subjected to sequential BLAST searches against the NCBI RefSeq viral nucleotide (blastn, 1e-5 E-value threshold) and protein (blastx, 1e-3) databases, and then the complete NCBI nucleotide (blastn, 1e-10) and protein (blastx, 1e-3) databases. In parallel, the adaptor and quality trimmed data were uploaded to Genome Detective (Vilsker et al. 2019), a web base software that assembles viral genomes from NGS data.

Bacteriophages, invertebrate viruses and retroviruses were excluded from subsequent steps, which only focused on eukaryotic, specifically vertebrate viruses. If our pipeline and Genome Detective generated a contig with the same viral hit, the contigs were compared. If they had the same sequence or were overlapping, the longest contig was selected. The filtered reads were mapped to the viral contigs to calculate the number of viral reads for each virus. Finally, the viral contigs were mapped to the reference genome of the virus corresponding to the best BLAST hit using Geneious v11.0.2 (Biomatters, Inc.). The baits were mapped to the same references to determine the genomic positions targeted by our bait panel for each virus.

### Phylogenetic analyses

Viral contigs were assigned to viral families according to the best BLAST results. Comprehensive sets of representative sequences from these viral families were retrieved from GenBank and aligned with the contigs using MAFFT v7.450 (Katoh & Standley 2013). Phylogenetic analysis was performed using the maximum-likelihood method based on the general time reversible substitution model with among-site rate heterogeneity modelled by the Γ distribution and estimation of proportion of invariable sites available in RAxML v8 (Stamatakis 2014), including 500 bootstrap replicates to determine node support. Phylogenetic analyses were performed only on viral contigs i) showing divergence from known viruses, i.e. with both BLAST identity and coverage to the best reference below 95%, to place them into a phylogenetic context, and ii) mapping to phylogenetically relevant genomic regions. Therefore, *Circoviridae* and *Anelloviridae* contigs were excluded as were those identified from water.

### Leech vertebrate host assignments

Host identification of leeches followed an eDNA/iDNA workflow (Axtner et al. 2019). In summary, leech samples were digested and short fragments of the mitochondrial markers 12S, 16S and cytochrome B were amplified in four PCR replicates each resulting in 12 PCR replicates per sample. We used a twin-tagging 2-step PCR protocol and PCR products were sequenced using an Illumina MiSeq (for details Axtner et al. 2019). After demultiplexing and read processing, each haplotype was taxonomically assigned to a curated reference database using PROTAX (Somervuo et al. 2016). Taxonomic assignments followed Axtner et al. (2019).

### Viral detection confirmation by PCR

The primers listed in Suppl. Tab. 2 were designed to confirm by PCR the viral contig sequences generated from the leech samples. For PCRs targeting RNA viruses, 50 uL of extract were digested with rDNase I (Ambion) following the manufacturer’s protocol. The DNAse-digested extracts were then purified using the RNeasy MinElute Cleanup Kit (Qiagen). RNA was reverse transcribed into cDNA using iScript^™^ Reverse Transcription Supermix (Bio Rad). Sediment and water samples that tested positive for EHV and JSRV were screened using a previously described pan-herpes PCR (Dayaram et al. 2017) and for JSRV (Palmarini et al. 2000), respectively. The resulting amplicons were Sanger sequenced.

### Shotgun sequencing

A subset of 64 leech samples were subjected to shotgun sequencing to evaluate the viral enrichment obtained by our capture system. An aliquot of pre-capture amplified genomic library of each leech sample was taken, pooled in equimolar amounts and sequenced on an Illumina MiSeq with the MiSeq Reagent Kit v3 (150-cycle). The shotgun reads were then mapped to the viral contigs obtained by viral capture using Bowtie2 v2.3.5.1 (Langmead & Salzberg 2012).

## RESULTS

### Positive and negative controls

In order to test the sensitivity of the viral capture in recovering vertebrate host viruses, the capture system was first applied to a positive control consisting of medical leeches fed with human blood spiked with two RNA viruses and two DNA viruses at different concentrations (Kampmann et al. 2017). All four viruses were detected, even if enrichment efficiency (proportion of on-target viral reads) and target genome recovery varied among viruses (Suppl. Fig. 1). No viral contigs were identified in the negative controls included to monitor laboratory contaminations for either the leech or water experiments. Further potential contamination from lab reagents was excluded (see Suppl. Information).

### Leech viral identification

Tiger leeches (*Haemadipsa picta*) and brown leeches (*Haemadipsa zeylanica*) were collected in Malaysian Borneo and processed as pools (bulk samples) consisting of 1 to 77 individual leeches separated by leech species and sampling location. Viruses were identified in 40 of the 68 leech pools analysed (59%) (Fig. 2; Suppl. Tab. 3). In 18 of these (45%), two to three viruses were identified. Sequence data from six vertebrate-infecting viral families were detected. The most common viral group was *Rhabdoviridae* found in 37% of samples, followed by *Coronaviridae* which was identified in 24% of samples. Members of the *Anelloviridae* were identified in 12% of samples, *Retroviridae* in 4%, and *Parvoviridae* and *Circoviridae* in 3%.

**Figure 2:**
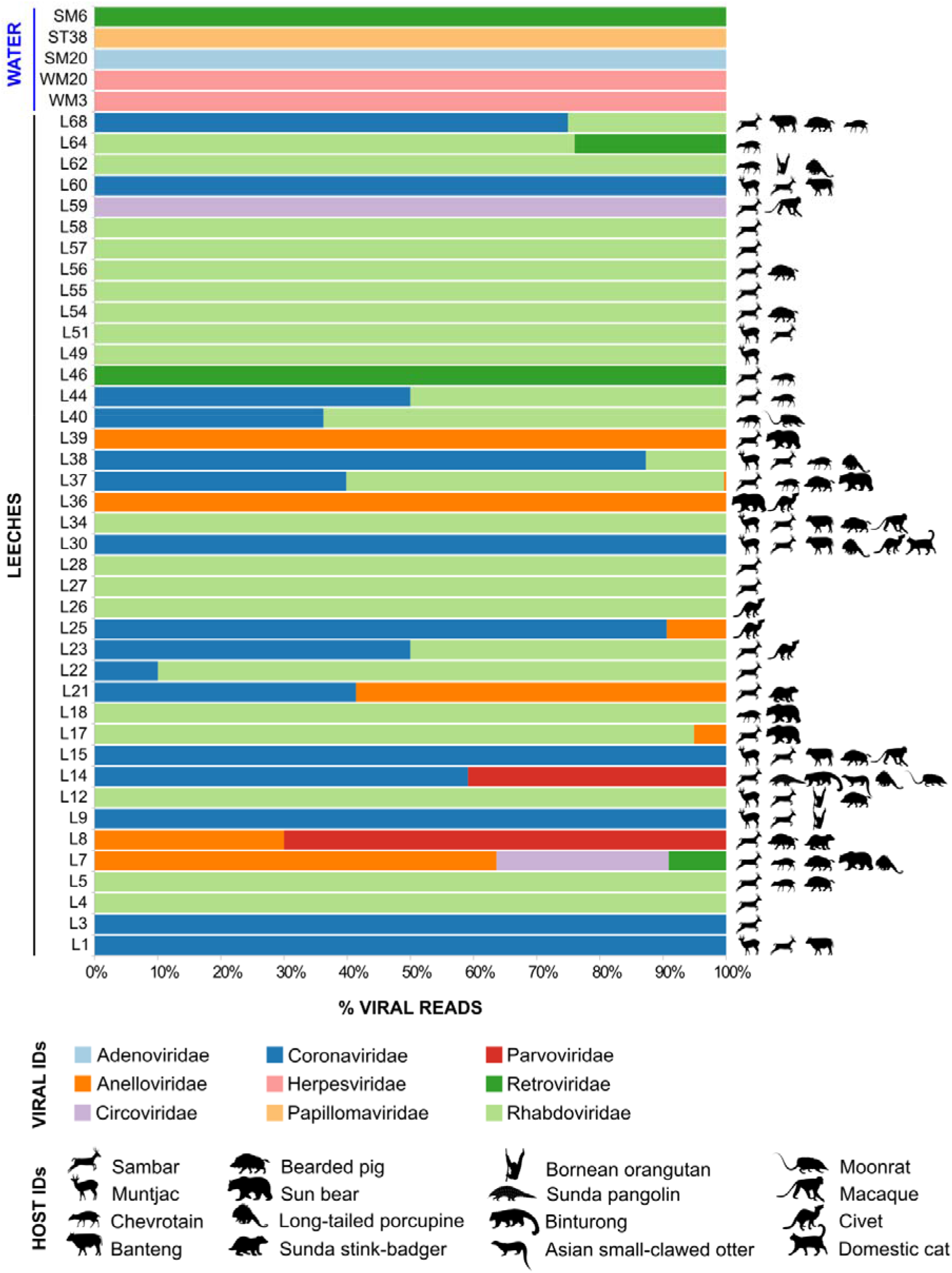
Relative abundance of viruses from each family, shown as the percentage of the total number of viral reads in each leech and waterhole sample. In the sample names, S stands for sediment, W for water, T for Tanzania, M for Mongolia and L for leeches. The leech host assignment for each leech sample is shown on the right (see Suppl. Tab. 3 for further details).

*Rhabdoviridae* contigs were genetically similar to three different viral genera (Suppl. Tab. 3). Five contigs were most similar (69-77%) to the Vesicular stomatitis Indiana virus (VSIV) (genus *Vesiculovirus*) as determined by BLAST searches. The limited similarity of these contigs to known rhabdoviruses suggest they may represent a new genus related to fish rhabodviruses (*Perhabdovirus* and *Sprivirus*) or *Vesiculovirus* (Suppl. Fig. 2). The other contigs clustered phylogenetically, suggesting they represent two new species of a rhabdovirus related to lyssaviruses (Suppl. Fig. 2). Although in most cases one contig per sample was observed, in five samples (L4, L12, L23, L58, L68) two different viruses were found. Most of the oligonucleotide baits were specific for the L gene which encodes the RNA-dependent RNA polymerase. All recovered contigs mapped to the L gene (Suppl. Fig. 3A-C). The viral contig sequences were confirmed by PCR and Sanger sequencing for L55 and L58 (Suppl. Fig. 3D).

All *Coronaviridae* contigs matched a bat betacoronavirus as determined by BLAST searches with identities between 70-73% (Suppl. Tab. 3). The resulting sequence did not cluster in any of the four clades representing the known *Coronaviridae* genera, suggesting it may represent a novel coronavirus genus (Suppl. Fig. 4). Each contig overlapped with the coronavirus *RNA-dependent RNA polymerase* gene (orf1ab), the viral region mainly targeted by the RNA oligonucleotide baits (Suppl. Fig. 5).

*Anelloviridae* contigs matched either porcine torque teno virus (PTTV) (95-96% identity), a giant panda anellovirus (GpAV) (81-92% identity) or a masked palm civet torque teno virus (Pl-TTV) (83-92% identity) (Suppl. Tab. 3). The PTTV contigs were found in two samples (L8 and L37), while the GpAV and Pl-TTV contigs were detected in six samples. GpAV was the best match in four samples (L7, L17, L36, L39) and Pl-TTV in three (L21, L25, L39). In sample L39 both were identified. Every *Anelloviridae* contig mapped to the non-coding region of the relative reference genome since all *Anelloviridae* baits targeted the same untranslated region (Suppl. Fig. 6A, C, E). The non-coding region sequenced is not phylogenetically informative and therefore, phylogenetic analysis could not be performed. Viral contigs were confirmed by PCR and Sanger sequencing for samples L7, L17, L25 and L37 (Suppl. Fig. 6B, D, F).

Three *Circoviridae* contigs matching a porcine circovirus (PCV) (100% identity) were identified in L7 and L59 (Suppl. Tab. 3). Two non-overlapping but adjacent contigs were retrieved from L7. A single contig overlapping with one of the two contigs determined from L7 was recovered from L59 (Suppl. Fig. 7A). The contigs mapped to the PCV replication protein (Rep), targeted by the *Circoviridae* baits (Suppl. Fig. 7A). The two overlapping contigs of L7 and L59 were confirmed by PCR and Sanger sequencing (Suppl. Fig. 7B). Since the identity of the contigs with known viral sequences in GenBank was 100%, no phylogenetic analysis was performed.

*Parvoviridae* contigs with the highest similarity to porcine parvovirus (PPV) were found in L8 (1 contig with 98% identity) and L14 (2 contigs with 74-77% identity) (Suppl. Tab. 3). The contig of L8 clustered within the *Tetraparvovirus* genus, close to ungulate parvoviruses (porcine, ovine and bovine PV), while the contigs of L14 within the *Copiparvovirus* genus, close to PPV4 (Suppl. Fig. 8). Two of the three contigs mapped to the *replicase* gene, while one from L14 mapped to an intergenic region (Suppl. Fig. 7C). Whereas the *replicase* region of PPV was covered by *Parvoviridae* baits, the intergenic region was not (Suppl. Fig. 7C). This portion of the virus may have been recovered by other non-*Parvoviridae* baits targeting that region non-specifically.

*Retroviridae* contigs similar to the simian and feline foamy virus (*Spumaretrovirinae* subfamily, 79-82% identity) were detected in three samples (L7, L46, L64) (Suppl. Tab. 3). Phylogenetically the contigs clustered together as a sister group to the feline foamy viruses (*Felispumavirus* genus), potentially being a new genus within the *Spumaretrovirinae* (Suppl. Fig. 9). The contigs mapped to the *polymerase* gene, which the exogenous retrovirus baits were designed to target (Suppl. Fig. 7D).

### Viral enrichment

Viral enrichment was low in both leech and waterholes samples, ranging from 0.00005% (sample L18) to 0.14% (sample L58) (average 0.01%), with a total of 12,710 viral reads out of 153,072,732 total reads (on-target proportion of 8,3E-5) found in 45 (40 leech and 5 waterhole samples) out of 89 total samples tested (68 leech and 21 waterhole samples) (Suppl. Tab. 4). We compared the results of capture of the leech samples to shotgun sequencing of a subset of 64 samples. In this subset, capture yielded 11,637 viral reads out of a total of 140,515,744 reads with an on-target proportion of 8,3E-5, whereas 4 viral reads were found by shotgun sequencing out of 4,961,274 total reads corresponding to an on-target proportion of 8,1E-7. In addition, the viral reads obtained by capture from leeches and waterholes were not randomly distributed across the viral genomes but aligned to the regions targeted by the baits (Suppl. Fig. 3, 5, 6, 7, 10, 11), reaching up to 479X (sample L58) mean depth of coverage in these regions (range 1.2X-479X). This and the two-fold order of magnitude increase in on-target viral reads, strongly indicates that enrichment of the viral genetic material present in the samples occurred.

### Leech bloodmeal host assignments

The mammalian hosts of the leeches were determined by metabarcoding (Axtner et al. 2019). The bearded pig (*Sus barbatus*) was identified in samples yielding porcine viruses, such as porcine circovirus (L7), porcine parvovirus (L8) and porcine torque teno virus (L8 and L37). Four leech samples with giant panda anellovirus (L7, L17, L36, L39) sequences yielded sun bear (*Helarctos malayanus*) sequences. Sequences aligning to the Malay civet (*Viverra tangalunga*) were identified in one of the three samples (L25) with masked palm civet torque teno virus sequences. Fourteen of the 16 samples with the potentially new coronavirus genus (87.5%) yielded deer sequences, specifically sambar (*Rusa unicolor*), indicating that the novel coronavirus might be a cervid virus. Similarly, the novel *Lyssavirus-like Rhabdoviridae* sequences were associated with cervid species (sambar or muntjac) (16 of 22 samples, 73%). However, due to the high prevalence of deer in the samples tested (70%) we could not reject that the occurrence of viruses and deer are independent variables (Chi^2^_Coronaviridae_ = 1.916, 1 df, p = 0.1663; Chi^2^_Rhabdoviridae_ = 1.046, 1 df, p = 0.3064).

### Waterhole viral identification

Five waterholes from Tanzania and six from Mongolia were tested. From each waterhole, one water filtrate and one sediment sample were collected (except for one Mongolia waterhole where only a sediment sample was collected), for a total of twenty-one samples. Five samples (two water and three sediment samples) were positive for viral sequences (23.8%). In filtered water and sediment samples collected from the same waterhole, only one virus per sample was identified and in one location (WM20 and SM20) contigs from different viral families were isolated based on sample type. Differences between sediment and water are not unexpected as the sediment likely represents a longer-term accumulation of biomaterial and the water represents more acute contamination at the surface and variable mixing throughout.

Of the 11 water filtrate samples tested, two samples from Mongolia (WM3 and WM20) (18.2%) had viral contigs with 100% identity to the Equid herpesvirus 1 and 3 (EHV-1 and EHV-3). The contig of WM20 mapped to the membrane glycoprotein B, whereas the two contigs of WM3 to the DNA packaging protein and membrane glycoprotein G, all regions covered by the *Herpesviridae* baits (Suppl. Fig. 10A-D). A nested panherpes PCR targeting the *DNA polymerase* gene and the resulting Sanger sequences further confirmed EHV presence (Suppl. Fig. 10E). Several equine species including domestic horses inhabit the Gobi Desert (Kaczensky et al. 2015), which is consistent with the presence of these viruses.

From the 12 sediment samples tested, two from Mongolia and one from Tanzania yielded viral sequences (25%) representing the *Retroviridae, Adenoviridae* and *Papillomaviridae* families. Mongolian sediment sample SM6 was positive for four contigs mapping to the *protease* (*pro*) gene of the Jaagsiekte sheep retrovirus (JSRV) with 100% identity (Suppl. Fig. 11A). JSRV from this sample was further confirmed by PCR (Suppl. Fig. 11A). Mongolia sediment sample SM20 was positive for Equine adenovirus (100% identity) with a contig mapping to a region comprised between the *pVI* and *hexon* capsid genes (Suppl. Fig. 11B). Given that multiple equine species are found in the Gobi Desert in Mongolia, it is likely that the water sources sampled may have been frequented by these species [10]. The sediment sample from Tanzania ST38 was positive for a Zetapapillomavirus related to the *Equus caballus* papillomavirus and *Equus asinus* papillomavirus (74% identity; *E1-E2* genes) (Suppl. Fig. 11C; Suppl. Fig. 12), consistent with the detection of Plains zebra’s (*Equus quagga*) DNA from this water source (Seeber et al. 2019).

## DISCUSSION

We demonstrate for the first time that both eDNA and iDNA sources can be used to survey known and novel viruses. Indeed, many of the viruses we identified from these two sources were highly divergent from available viral reference genomes (homology 45-100%, average 80%). We show that DNA and RNA viruses could be detected in 59% and 23.8% of the iDNA (leech) and eDNA (waterhole) samples, respectively. The congruence of host DNA assignment for leeches and viral families identified suggests that bloodmeals are useful for determining viral diversity. The detection of equine viruses from African and Mongolian waterholes, where intense wild equid visitation rates were directly observed, suggests eDNA derived viruses reflect host utilization of the water rather than other environmental sources such as fomites. While host assignments are difficult to establish for novel viruses from eDNA and to a lesser extent in iDNA, as e/iDNA samples can contain DNA of multiple hosts, the results do narrow the possible number of taxa down to a small portion of the overall faunal diversity within the regions examined. For example, the novel coronavirus was strongly, but not in a statistically significant way, associated with sambar. This suggests targeted sampling and virus specific PCRs could be used to examine prevalence in the species and to establish whether they are a potential viral reservoir. Narrowing down the potential taxa that need to be screened in biodiversity hotspots will be critical, particularly for viruses hosted at low prevalence within hotspot regions.

PCR based approaches have been used to detect known pathogens from flies (Bitome-Essono et al. 2017; Gogarten et al. 2019) or from medicinal leeches under laboratory conditions (Kampmann et al. 2017). While major findings have resulted from such analysis, PCR based approaches are often poorly suited to the discovery of novel viruses that may be highly divergent as PCR based approaches often fail when viral divergence exceeds 5-10%, particularly relevant to RNA viruses (Schlaberg et al. 2017). The unknown viral diversity in the wild, and the potential degradation of viral nucleic acids in bloodmeals or in the environment, may affect PCR detection resulting in high false negative rates. Hybridization capture overcomes such limitations because the short baits can capture divergent and degraded DNA. With our capture system we were able to identify viral sequences with up to 55% divergence from known viral genomes. The comprehensive viral group representation in the bait set allows for the determination of both viral presence and viral diversity. The ability of oligonucleotides with substantial divergence from the target sequence to capture more distantly related sequences is particularly useful in virology since most viruses are uncharacterized in wildlife and many evolve rapidly (Howard & Fletcher 2012). The overall enrichment was low, but this was not unexpected since it is unlikely that any leech sample was strongly viremic or that large amount of virus was shed into water. Nevertheless, our viral capture system generated a two fold order of magnitude higher viral enrichment compared to shotgun sequencing. Furthermore, viral enrichment was concentrated in the regions of the viral genome where baits were designed, leading to high coverage (up to 479X) at these positions. This allowed for the assembly of viral genome fragments for which we could reconstruct the phylogenies and enabled further PCR methods to be successfully implemented to confirm some of the viral capture results.

Using short RNA baits to capture highly conserved sequences from every known vertebrate viral genome is a useful and relatively inexpensive approach for providing an initial viral identification (Figure 1). However, to fully characterize each virus, the RNA oligonucleotide bait set would need to be customized to retrieve full length viral genomes which has also been done successfully for novel divergent viruses (Alfano et al. 2016). This is a possible strategy to further investigate only viruses of interest, whereas initial screening with full length genomes for all viruses is likely too costly.

Several novel viruses were identified from leech bloodmeals with our approach, which is not unexpected as little is known about the virology of wildlife in Southeast Asia, where the leeches were collected. Several viral contigs were phylogenetically distinct from known viruses and may represent new genera. For example, the novel coronavirus identified in leech bloodmeals did not cluster with any of the known *Coronaviridae* clades. We could also tentatively associate the novel corona- and rhabdoviruses with cervids, which are regularly sold as bushmeat in wildlife markets (Nasi et al. 2011). Both recent coronavirus epidemics (SARS-CoV (Drosten et al. 2003) and SARS-Cov-2 (Zhou et al. 2020) spilled over from wildlife. This suggests that e/iDNA-based pathogen surveillance approaches may complement efforts to proactively identify novel viruses that could potentially spillover to humans or livestock.

The collection of wild haematophagous invertebrates such as leeches or water and sediments has both advantages and disadvantages compared to invasively collected wildlife samples. Large amounts of DNA can be extracted from bloodmeals, in particular when leeches are processed in bulk. We pooled up to 77 leeches and many of our leech bulk samples contained a diverse mix of mammalian DNA. A disadvantage of leeches is that they cannot be found in all environments: for example haematophagous terrestrial species are restricted to tropical rainforests of Asia, Madagascar and Australia (Schnell et al. 2018). In addition, leech feeding biases could influence diversity surveys (Abrams et al. 2019; Schnell et al. 2015). However, this disadvantage could be overcome in the future by employing additional invertebrates such mosquitoes (Ng et al. 2011) or carrion flies (Hoffmann et al. 2016). Waterholes are commonly found in almost all environments. In environments with seasonal water shortages, DNA from animals can become highly concentrated due to many animals utilizing rare water sources. The disadvantages are that the dilution factor of water, depending on water body size, can obscure rare DNA sequences and mixed host species sequences are generally the rule rather than the exception. Further experiments with field filtration and sample concentration such as methods used with pathogen detection in waste water may improve detection rates (Farkas et al. 2018).

Environmental DNA and in particular its subdiscipline invertebrate-derived DNA viral hybridization capture may be a useful and economical tool for viral identification and characterization particularly in difficult to access sampling environments prior to potential viral emergence. Sampling in environments where direct access to animals is difficult or highly restricted, eDNA and iDNA may be the only option to detect viruses circulating in the wild. Paired with fecal samples and iDNA from other invertebrates would greatly broaden the substrates from which viral surveillance could be performed. The current study suggests this approach will be successful in complementing invasive approaches or replacing them in environments where invasive approaches are not possible.

## ACKNOWLEDGEMENTS

We would like to thank Joseph DeRisi for guidance in using his oligonucleotide data. We thank Peter Seeber and Sanatana-Erini Soilemetzidou for collection of water and sediment samples in Africa and Mongolia respectively. We thank the Sabah Forestry Department, especially Johnny Kissing, Peter Lagan, and Datuk Sam Mannan, for supporting the fieldwork and the Sabah Biodiversity Council for providing research, collection, and export permits (JKM/MBS.1000-2/3 JLD.2) for the leech work. This project received financial support from the German Federal Ministry of Education and Research (BMBF FKZ: 01LN1301A) to NA, JA, AM and AW and was supported by funds from the Leibniz Gemeinschaft, SAW-2015-IZW-1 440 to AD and ADG.

## AUTHORS’ CONTRIBUTIONS

ADG and AW designed the study; AM and STW collected the leeches in the field; JA performed the leech nucleic acids extractions and the PCRs on leeches; NA and AD performed the capture experiments; NA and KT performed the bioinformatics and phylogenetic analyses; NA, AD, ADG and AW wrote the manuscript; NA, AD, JA, KT, MLK, TPG, AW and ADG reviewed the manuscript; all authors read and approved the final manuscript.

## DATA AVAILABILITY

The dataset generated and analyzed during this study is available in the NCBI Sequence Read Archive (SRA) repository under the accession number PRJNA627811. The consensus sequences of the viral contigs obtained in this study were deposited in GenBank under the accession numbers MT444451-MT444487.

## SUPPORTING INFORMATION

Supporting information is available at MEE online.

## REFERENCES

Abrams, J. F., Hörig, L. A., Brozovic, R., Axtner, J., Crampton-Platt, A., Mohamed, A., Wong, S. T., et al. (2019). ‘Shifting up a gear with iDNA: From mammal detection events to standardised surveys’, Journal of Applied Ecology, 56/7: 1637–1648.

Alfano, Niccoló, Courtiol, A., Vielgrader, H., Timms, P., Roca, A. L., & Greenwood, A. D. (2015). ‘Variation in koala microbiomes within and between individuals: effect of body region and captivity status’, Scientific reports, 5: 10189.

Alfano, Niccolò, Michaux, J., Morand, S., Aplin, K., Tsangaras, K., Löber, U., Fabre, P.-H., et al. (2016). ‘Endogenous gibbon ape leukemia virus identified in a rodent (Melomys burtoni subsp.) from Wallacea (Indonesia)’, Journal of virology, 90/18: 8169–8180. Am Soc Microbiol.

Axtner, J., Crampton-Platt, A., Hörig, L. A., Mohamed, A., Xu, C. C., Yu, D. W., & Wilting, A. (2019). ‘An efficient and robust laboratory workflow and tetrapod database for larger scale environmental DNA studies’, GigaScience, 8/4: giz029.

Bankevich, A., Nurk, S., Antipov, D., Gurevich, A. A., Dvorkin, M., Kulikov, A. S., Lesin, V. M., et al. (2012). ‘SPAdes: a new genome assembly algorithm and its applications to single-cell sequencing’, Journal of computational biology, 19/5: 455–477.

Bitome-Essono, P.-Y., Ollomo, B., Arnathau, C., Durand, P., Mokoudoum, N. D., Yacka-Mouele, L., Okouga, A.-P., et al. (2017). ‘Tracking zoonotic pathogens using blood-sucking flies as’ flying syringes’’, elife, 6: e22069.

Chen, E. C., Miller, S. A., DeRisi, J. L., & Chiu, C. Y. (2011). ‘Using a Pan-Viral Microarray Assay (Virochip) to Screen Clinical Samples for Viral Pathogens’,.

Daszak, P., Cunningham, A. A., & Hyatt, A. D. (2000). ‘Emerging infectious diseases of wildlife–threats to biodiversity and human health’, science, 287/5452: 443–449. American Association for the Advancement of Science.

Dayaram, A., Franz, M., Schattschneider, A., Damiani, A. M., Bischofberger, S., Osterrieder, N., & Greenwood, A. D. (2017). ‘Long term stability and infectivity of herpesviruses in water’, Scientific reports, 7: 46559.

Drosten, C., Günther, S., Preiser, W., Van Der Werf, S., Brodt, H.-R., Becker, S., Rabenau, H., et al. (2003). ‘Identification of a novel coronavirus in patients with severe acute respiratory syndrome’, New England journal of medicine, 348/20: 1967–1976.

Edgar, R. C. (2010). ‘Search and clustering orders of magnitude faster than BLAST’, Bioinformatics, 26/19: 2460–2461.

Farkas, K., McDonald, J. E., Malham, S. K., & Jones, D. L. (2018). ‘Two-step concentration of complex water samples for the detection of viruses’, Methods and protocols, 1/3: 35.

Gogarten, J. F., Düx, A., Mubemba, B., Pléh, K., Hoffmann, C., Mielke, A., Müller-Tiburtius, J., et al. (2019). ‘Tropical rainforest flies carrying pathogens form stable associations with social nonhuman primates’, Molecular ecology, 28/18: 4242–4258.

Grabherr, M. G., Haas, B. J., Yassour, M., Levin, J. Z., Thompson, D. A., Amit, I., Adiconis, X., et al. (2011). ‘Trinity: reconstructing a full-length transcriptome without a genome from RNA-Seq data’, Nature biotechnology, 29/7: 644.

Hoffmann, C., Stockhausen, M., Merkel, K., Calvignac-Spencer, S., & Leendertz, F. H. (2016). ‘Assessing the feasibility of fly based surveillance of wildlife infectious diseases’, Scientific reports, 6/1: 1–9.

Howard, C. R., & Fletcher, N. F. (2012). ‘Emerging virus diseases: can we ever expect the unexpected?’, Emerging microbes & infections, 1/1: 1–9. Taylor & Francis.

Hoye, B. J., Munster, V. J., Nishiura, H., Klaassen, M., & Fouchier, R. A. (2010). ‘Surveillance of wild birds for avian influenza virus’, Emerging infectious diseases, 16/12: 1827.

Johnson, J., Howard, K., Wilson, A., Ward, M., Gilbert, G. L., & Degeling, C. (2019). ‘Public preferences for One Health approaches to emerging infectious diseases: a discrete choice experiment’, Social Science & Medicine, 228: 164–171.

Jones, K. E., Patel, N. G., Levy, M. A., Storeygard, A., Balk, D., Gittleman, J. L., & Daszak, P. (2008). ‘Global trends in emerging infectious diseases’, Nature, 451/7181: 990–993.

Kaczensky, P., Lkhagvasuren, B., Pereladova, O., Hemami, M., & Bouskila, A. (2015). Equus hemionus. The IUCN Red List of Threatened Species 2015: e. T7951A45171204.

Kampmann, M.-L., Schnell, I. B., Jensen, R. H., Axtner, J., Sander, A. F., Hansen, A. J., Bertelsen, M. F., et al. (2017). ‘Leeches as a source of mammalian viral DNA and RNA—a study in medicinal leeches’, European journal of wildlife research, 63/2: 36.

Katoh, K., & Standley, D. M. (2013). ‘MAFFT multiple sequence alignment software version 7: improvements in performance and usability’, Molecular biology and evolution, 30/4: 772–780. Society for Molecular Biology and Evolution.

Kopylova, E., Noé, L., & Touzet, H. (2012). ‘SortMeRNA: fast and accurate filtering of ribosomal RNAs in metatranscriptomic data’, Bioinformatics, 28/24: 3211–3217.

Langmead, B., & Salzberg, S. (2012). Fast gapped-read alignment with bowtie 2 Nat Methods 9 (4): 357–359. pmid: 22388286 View Article PubMed. NCBI.

Leroy, E. M., Epelboin, A., Mondonge, V., Pourrut, X., Gonzalez, J.-P., Muyembe-Tamfum, J.-J., & Formenty, P. (2009). ‘Human Ebola outbreak resulting from direct exposure to fruit bats in Luebo, Democratic Republic of Congo, 2007’, Vector-borne and zoonotic diseases, 9/6: 723–728. Mary Ann Liebert, Inc. 140 Huguenot Street, 3rd Floor New Rochelle, NY 10801 USA.

Meyer, M., & Kircher, M. (2010). ‘Illumina sequencing library preparation for highly multiplexed target capture and sequencing’, Cold Spring Harbor Protocols, 2010/6: pdb–prot5448.

Mosher, B. A., Huyvaert, K. P., Chestnut, T., Kerby, J. L., Madison, J. D., & Bailey, L. L. (2017). ‘Design-and model-based recommendations for detecting and quantifying an amphibian pathogen in environmental samples’, Ecology and evolution, 7/24: 10952–10962.

Nasi, R., Taber, A., & Van Vliet, N. (2011). ‘Empty forests, empty stomachs? Bushmeat and livelihoods in the Congo and Amazon Basins’, International Forestry Review, 13/3: 355–368.

Ng, T. F. F., Willner, D. L., Lim, Y. W., Schmieder, R., Chau, B., Nilsson, C., Anthony, S., et al. (2011). ‘Broad surveys of DNA viral diversity obtained through viral metagenomics of mosquitoes’, PloS one, 6/6.

Palmarini, M., Datta, S., Omid, R., Murgia, C., & Fan, H. (2000). ‘The long terminal repeat of Jaagsiekte sheep retrovirus is preferentially active in differentiated epithelial cells of the lungs’, Journal of virology, 74/13: 5776–5787. Am Soc Microbiol.

Pruvot, M., Khammavong, K., Milavong, P., Philavong, C., Reinharz, D., Mayxay, M., Rattanavong, S., et al. (2019). ‘Toward a quantification of risks at the nexus of conservation and health: The case of bushmeat markets in Lao PDR’, Science of the total environment, 676: 732–745.

Schlaberg, R., Queen, K., Simmon, K., Tardif, K., Stockmann, C., Flygare, S., Kennedy, B., et al. (2017). ‘Viral pathogen detection by metagenomics and pan-viral group polymerase chain reaction in children with pneumonia lacking identifiable etiology’, The Journal of infectious diseases, 215/9: 1407–1415. Oxford University Press.

Schnell, I. B., Bohmann, K., Schultze, S. E., Richter, S. R., Murray, D. C., Sinding, M.-H. S., Bass, D., et al. (2018). ‘Debugging diversity–a pan-continental exploration of the potential of terrestrial blood-feeding leeches as a vertebrate monitoring tool’, Molecular ecology resources, 18/6: 1282–1298.

Schnell, I. B., Sollmann, R., Calvignac-Spencer, S., Siddall, M. E., Douglas, W. Y., Wilting, A., & Gilbert, M. T. P. (2015). ‘iDNA from terrestrial haematophagous leeches as a wildlife surveying and monitoring tool–prospects, pitfalls and avenues to be developed’, Frontiers in zoology, 12/1: 24.

Seeber, P. A., McEwen, G. K., Löber, U., Förster, D. W., East, M. L., Melzheimer, J., & Greenwood, A. D. (2019). ‘Terrestrial mammal surveillance using hybridization capture of environmental DNA from African waterholes’, Molecular ecology resources.

Sharp, P. M., & Hahn, B. H. (2011). ‘Origins of HIV and the AIDS pandemic’, Cold Spring Harbor perspectives in medicine, 1/1: a006841.

Śmietanka, K., Woźniakowski, G., Kozak, E., Niemczuk, K., Frączyk, M., Bocian, \Lukasz, Kowalczyk, A., et al. (2016). ‘African swine fever epidemic, Poland, 2014–2015’, Emerging infectious diseases, 22/7: 1201.

Somervuo, P., Koskela, S., Pennanen, J., Henrik Nilsson, R., & Ovaskainen, O. (2016). ‘Unbiased probabilistic taxonomic classification for DNA barcoding’, Bioinformatics, 32/19: 2920–2927.

Stamatakis, A. (2014). ‘RAxML version 8: a tool for phylogenetic analysis and post-analysis of large phylogenies’, Bioinformatics, 30/9: 1312–1313.

Swift, L., Hunter, P. R., Lees, A. C., & Bell, D. J. (2007). ‘Wildlife trade and the emergence of infectious diseases’, EcoHealth, 4/1: 25. Springer.

Tilker, A., Abrams, J. F., Nguyen, A., Hörig, L., Axtner, J., Louvrier, J., Rawson, B. M., et al. (2020). ‘Identifying conservation priorities in a defaunated tropical biodiversity hotspot’, Diversity and Distributions, 26/4: 426–440. Wiley Online Library.

Vilsker, M., Moosa, Y., Nooij, S., Fonseca, V., Ghysens, Y., Dumon, K., Pauwels, R., et al. (2019). ‘Genome Detective: an automated system for virus identification from high-throughput sequencing data’, Bioinformatics, 35/5: 871–873.

Wang, D., Coscoy, L., Zylberberg, M., Avila, P. C., Boushey, H. A., Ganem, D., & DeRisi, J. L. (2002). ‘Microarray-based detection and genotyping of viral pathogens’, Proceedings of the National Academy of Sciences, 99/24: 15687–15692.

Wilking, H., Ziller, M., Staubach, C., Globig, A., Harder, T. C., & Conraths, F. J. (2009). ‘Chances and limitations of wild bird monitoring for the avian influenza virus H5N1—detection of pathogens highly mobile in time and space’, PLoS One, 4/8.

Yozwiak, N. L., Skewes-Cox, P., Stenglein, M. D., Balmaseda, A., Harris, E., & DeRisi, J. L. (2012). ‘Virus identification in unknown tropical febrile illness cases using deep sequencing’, Plos neglected tropical diseases, 6/2.

Zhou, P., Yang, X.-L., Wang, X.-G., Hu, B., Zhang, L., Zhang, W., Si, H.-R., et al. (2020). ‘A pneumonia outbreak associated with a new coronavirus of probable bat origin’, Nature, 1–4.

