## supplemental material for "Non-invasive surveys of mammalian viruses using environmental DNA"

### Supporting Information

#### Suid DNA and contamination

It is known to that lab reagents are often contaminated by DNA from various species including pig (*Sus scrofa*) (Leonard et al. 2007). We identified porcine viruses, such as porcine circovirus, porcine parvovirus and, torque teno sus virus in our samples. The fact the we did not find these sequences in the negative controls argues against general pig DNA contamination. Moreover, Sus barbatus, an endemic pig species found in the region where leeches were sampled, was identified by metabarcoding (Axtner et al. 2019) as leech host in 3 (L7, L8, L37) out the 5 samples positive to porcine viruses. Furthermore, in order to understand if there was pig DNA contamination in our data, we mapped the reads from the samples which were positive for porcine viruses to the mitochondrial genomes of both Sus scrofa and Sus barbatus, since often mitochondrial genomes are obtained from capture experiments as byproducts, and we observed that the mitochondrial reads had higher identity to the Sus barbatus mitochondrial genome than to Sus scrofa. Finally, pig viruses were not identified in the water samples. This suggests that the porcine viruses found in the leech samples did not derive from pig, *Sus scrofa*, DNA contamination, but rather from bearded pigs the leech had fed on.

#### Supplementary Figure Legends

**Suppl. Fig. 1.** Application of viral capture on medical leeches fed with human blood spiked with four viruses, as positive control. All four viruses were detected, but enrichment efficiency (proportion of on-target viral reads) (bottom left) and target genome recovery (bottom right) varied among viruses. At the bottom right, the contigs recovered for each virus (black bars) are mapped to the reference genome together with the corresponding RNA olignonucleotide baits (blue bars). The adeno-associated virus (AAV) recovered by capture may have been present in the HeLa cell line in which HAdV was cultivated.

**Suppl. Fig. 2.** Evolutionary relationships of the partial *L* gene sequence of the rhabdoviruses identified in the leech bloodmeals (in red) with representatives of the *Rhabdoviridae* family. The consensus sequences of the contigs found in leech samples were used as representative sequence. The main genera of the *Rhabdoviridae* family are indicated on the right. The tree was inferred using the maximum-likelihood method with the GTR+G+I model and node robustness assessed with 500 rapid bootstrap pseudoreplicates. The tree is scaled according to the number of nucleotide substitutions per site. The tree is rooted arbitrarily on one of two basal clades (genera *Almendravirus* and *Bahiavirus*) that comprise viruses isolated from mosquitoes. Abbreviations and accession numbers of reference sequences are reported in https://doi.org/10.1371/journal.ppat.1004664.s013 (Walker et al. 2015).

**Suppl. Fig. 3.** Genomic regions covered by the *Vesiculovirus* (A), *Ephemerovirus* (B) and *Lyssavirus* (C) contigs (black bars) identified in leech bloodmeals and corresponding RNA oligonucleotides bait (blue bars) positions. The sequences of the PCR primers and PCR products used to confirm the *Lyssavirus* contigs are shown in panel D.

**Suppl. Fig. 4.** Evolutionary relationships of the partial *RNA-dependent RNA polymerase* gene (orf1ab) sequence of the coronavirus identified in the leech bloodmeals (in red) with representatives of the *Coronaviridae* family. The consensus of the 16 contigs found in leech samples was used as representative sequence. The four genera (*Alpha*-, *Beta*-, *Gamma*- and *Delta-coronavirus*) of the *Coronaviridae* family are indicated. The betacoronaviruses are further divided into four (a-d) subgroup clusters. The tree was inferred using the maximum-likelihood method with the GTR+G+I model and node robustness assessed with 500 rapid bootstrap pseudoreplicates. The tree is scaled according to the number of nucleotide substitutions per site. The tree is midpoint-rooted for purposes of clarity. CoV: coronavirus.

**Suppl. Fig. 5.** Genomic region covered by the *Coronaviridae* contigs (black bars) identified in leech bloodmeals and corresponding RNA oligonucleotides bait (blue bars) positions.

**Suppl. Fig. 6.** Genomic regions covered by contigs (black bars) of the three *Anelloviridae* – Torque teno sus virus (A), giant panda anellovirus (C) and *Paguma larvata* torque teno virus (E) – identified in leech bloodmeals and corresponding RNA oligonucleotides bait (blue bars) positions. The sequences of the PCR primers and PCR products used to confirm the Torque teno sus virus, giant panda anellovirus and *Paguma larvata* torque teno virus contigs are shown in panels B, D and F, respectively.

**Suppl. Fig. 7.** Genomic regions covered by the porcine circovirus (A), porcine parvovirus (C) and feline foamy virus (D) contigs (black bars) identified in leech bloodmeals and corresponding RNA oligonucleotides bait (blue bars) positions. The sequences of the PCR primers and PCR products used to confirm the porcine circovirus contigs are shown in panels B.

**Suppl. Fig. 8.** Evolutionary relationships of the partial *replicase* (*NS1*) gene sequence of the parvovirus identified in the leech bloodmeals (in red) with representatives of the *Parvovirinae* subfamily (*Parvoviridae* family). The main genera (*Ave*-, *Copi*-, *Dependo*-, *Erythro*-, *Proto*-, and *Tetra-parvovirus*) of the *Parvovirinae* subfamily are indicated on the right. The tree was inferred using the maximum-likelihood method with the GTR+G+I model and node robustness assessed with 500 rapid bootstrap pseudoreplicates. The tree is scaled according to the number of nucleotide substitutions per site. The tree is midpoint-rooted for purposes of clarity. PV: parvovirus; AAV: adeno-associated virus.

**Suppl. Fig. 9.** Evolutionary relationships of the partial *polymerase* gene sequence of the foamy viruses identified in the leech bloodmeals (in red) with representatives of the *Spumaretrovirinae* subfamily (family *Retroviridae*). The four genera (*Simii*-, *Equi*-, *Bovi*- and *Feli-spumavirus*) of the *Spumaretrovirinae* subfamily are indicated on the right. The tree was inferred using the maximum-likelihood method with the GTR+G+I model and node robustness assessed with 500 rapid bootstrap pseudoreplicates. The tree is scaled according to the number of nucleotide substitutions per site. The tree is rooted on the clade of the sloth endogenous foamy viruses (SloEFV). FV: foamy virus.

**Suppl. Fig. 10.** Genomic regions covered by the Equid herpesvirus (EHV) contigs (red bars) identified in the water samples and corresponding RNA oligonucleotides bait (blue bars) positions. An enlargement of the three different regions covered is shown in panels B, C and D. The sequence of the product of the nested panherpes PCR targeting the *DNA polymerase* gene which was used to confirm the presence of EHV in the samples is shown in panels A and E (enlargement) in green.

**Suppl. Fig. 11.** Genomic regions covered by the *Retroviridae* (A), *Adenoviridae* (B) and *Papillomaviridae* (C) contigs (black bars) identified in sediment samples and corresponding RNA oligonucleotides bait (blue bars) positions. The sequence of the product of the PCR targeting the *env* gene which was used to confirm the presence of JSRV in sample SM3 is shown in light green (A).

**Suppl. Fig. 12.** Evolutionary relationships of the partial *E1-E2* gene sequence of the papillomavirus identified in a sediment sample (in red) with representatives of the *Papillomaviridae* family. The main genera of the *Papillomaviridae* family are indicated on the right. The tree was inferred using the maximum-likelihood method with the GTR+G+I model and node robustness assessed with 500 rapid bootstrap pseudoreplicates. The tree is scaled according to the number of nucleotide substitutions per site. The tree is midpoint-rooted for purposes of clarity. PV: papillomavirus.

#### Supplementary References

Axtner, J., Crampton-Platt, A., Hörig, L. A., Mohamed, A., Xu, C. C., Yu, D. W., & Wilting, A. (2019). 'An efficient and robust laboratory workflow and tetrapod database for larger scale environmental DNA studies', *GigaScience*, 8/4: giz029.

Leonard, J. A., Shanks, O., Hofreiter, M., Kreuz, E., Hodges, L., Ream, W., Wayne, R. K., et al. (2007). 'Animal DNA in PCR reagents plagues ancient DNA research', *Journal of Archaeological Science*, 34/9: 1361–6. DOI: https://doi.org/10.1016/j.jas.2006.10.023

S1. Walker, P.J., Firth, C., Widen, S.G., Blasdell, K.R., Guzman, H., et al. (2015). *Evolution of Genome Size and Complexity in the Rhabdoviridae. PLOS Pathogens, 11(2): e1004664.*

#### Supplementary Table Legends

**Suppl. Tab. 1.** Read counts for leech and waterhole samples with SRA accession numbers.

**Suppl. Tab. 2.** Primers used and amplification results obtained for the leech viral contigs confirmation PCRs.

**Suppl. Tab. 3.** Best BLAST hit for each viral contig identified in leech and waterhole samples.

**Suppl. Tab. 4.** Viral enrichment achieved by capture for each sample.

**Suppl. Fig. 1**

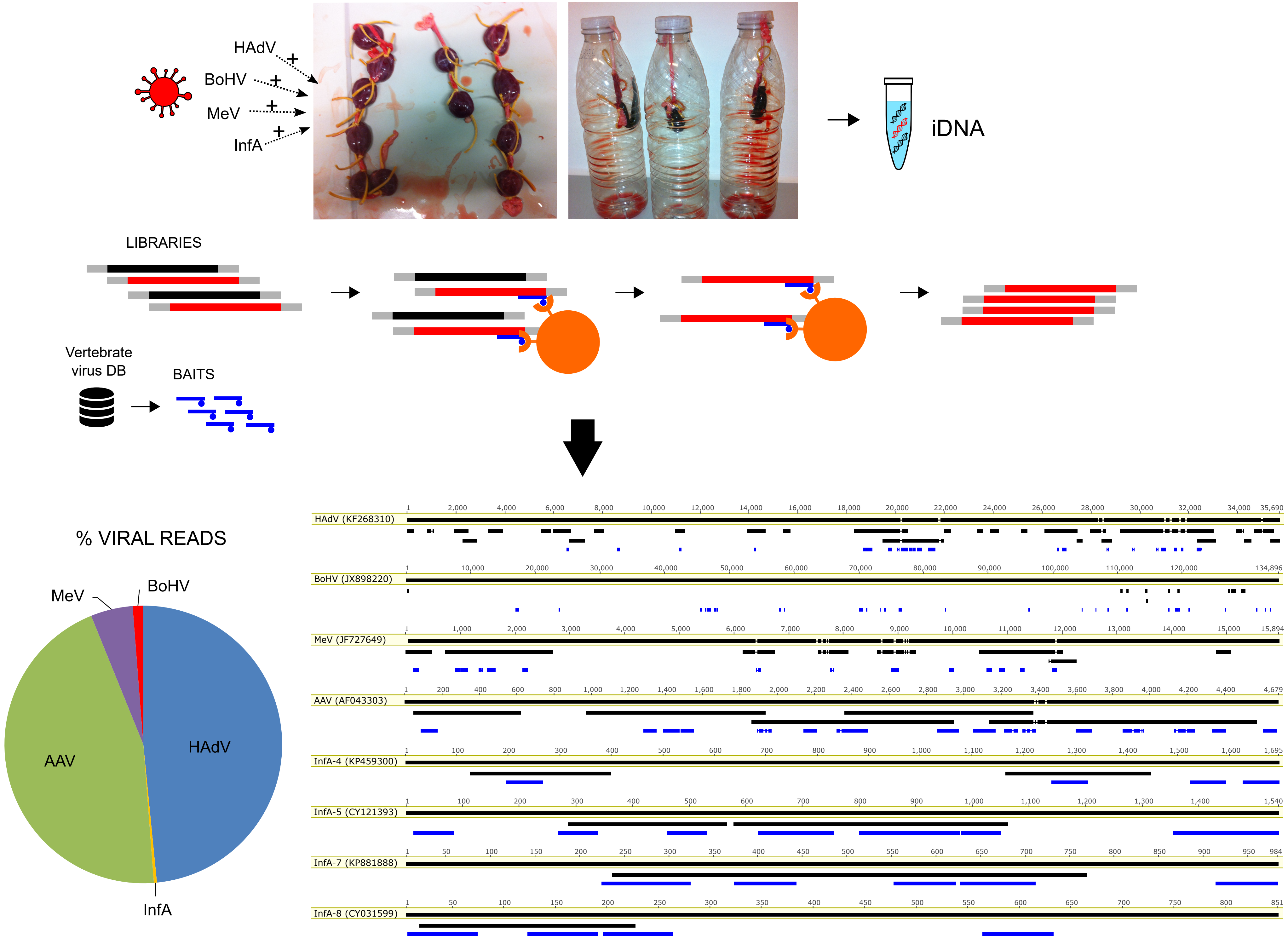
HAdV
BoHV
MeV
InfA
iDNA
LIBRARIES
Vertebrate virus DB
BAITS
% VIRAL READS
MeV
BoHV
AAV
HAdV
InfA
HAdV (KF268310)
BoHV (JX898220)
MeV (JF727649)
AAV (AF043303)
InfA-4 (KP459300)
InfA-5 (CY121393)
InfA-7 (KP881888)
InfA-8 (CY031599)

**Suppl. Fig. 2**

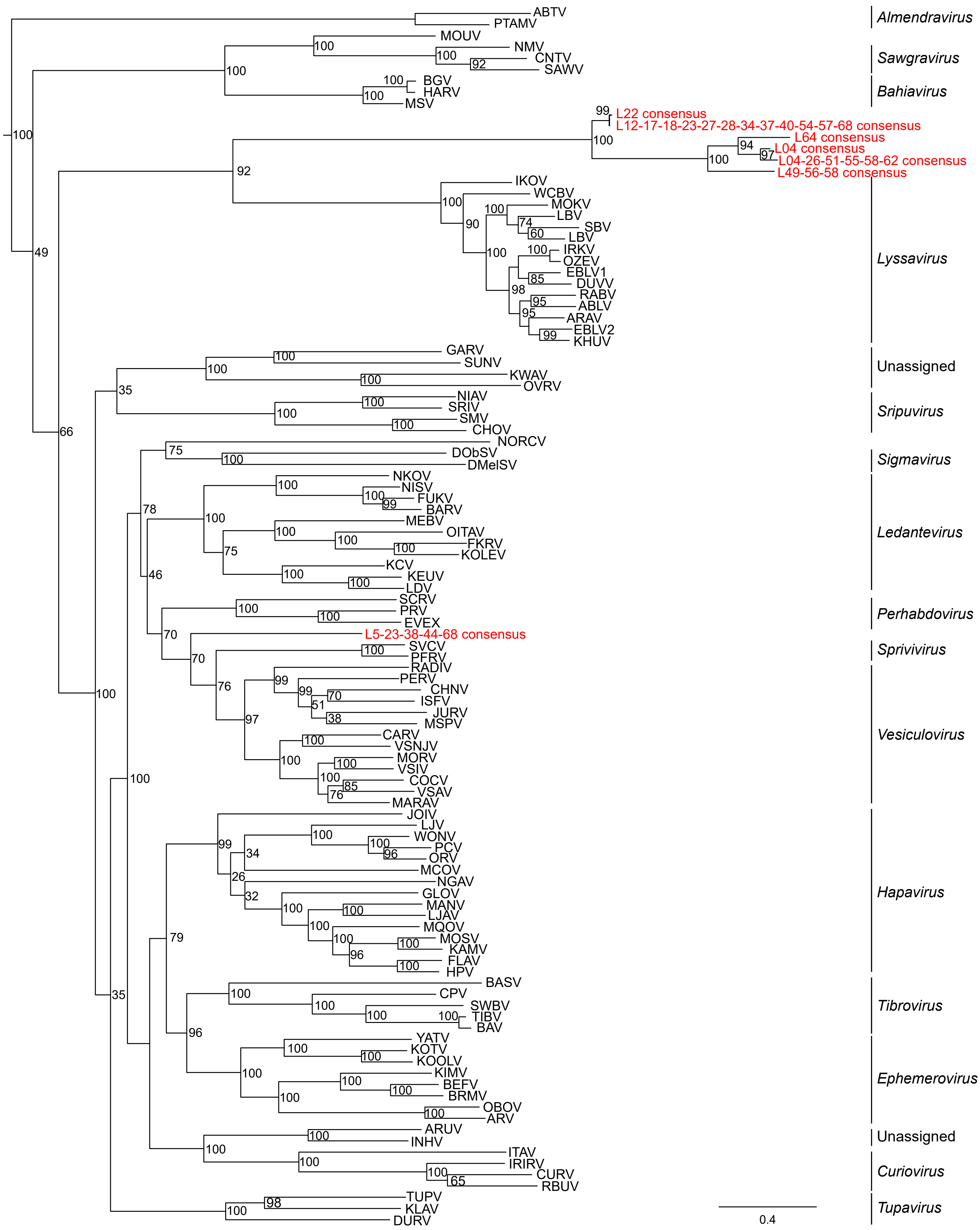

ABTV
PTAMV
Almendravirus
MOUV
NMV
CNTV
SAWV
Sawgravirus
BGV
HARV
MSV
Bahiavirus
L22 consensus
L12-17-18-23-27-28-34-37-40-54-57-68 consensus
L64 consensus
L04 consensus
L04-26-51-55-58-62 consensus
L49-56-58 consensus
IKOV
WCBV
MOKV
LBV
SBV
LBV
IRKV
OZEV
EBLV1
DUVV
RABV
ABLV
ARAV
EBLV2
KHUV
Lyssavirus
GARV
SUNV
KWAV
OVRV
Unassigned
NIAV
SRIV
SMV
CHOV
Sripuvirus
NORCV
DObSV
DMelSV
Sigmavirus
NKOV
NISV
FUKV
BARV
MEBV
OITAV
FKRV
KOLEV
KCV
KEUV
LDV
Ledantevirus
SCRV
PRV
EVEX
Perhabdovirus
L5-23-38-44-68 consensus
SVCV
PFRV
Sprivivirus
RADIV
PERV
CHNV
ISFV
JURV
MSPV
CARV
VSNJV
MORV
VSIV
COCV
VSAV
MARAV
Vesiculovirus
JOIV
LJV
WONV
PCV
ORV
MCOV
NGAV
GLOV
MANV
LJAV
MQOV
MOSV
KAMV
FLAV
HPV
Hapavirus
BASV
CPV
SWBV
TIBV
BAV
Tibrovirus
YATV
KOTV
KOOLV
KIMV
BEFV
BRMV
OBOV
ARV
Ephemerovirus
ARUV
INHV
Unassigned
ITAV
IRIRV
CURV
RBUV
Curiovirus
TUPV
KLAV
DURV
Tupavirus
0.4

#### Suppl. Fig. 3

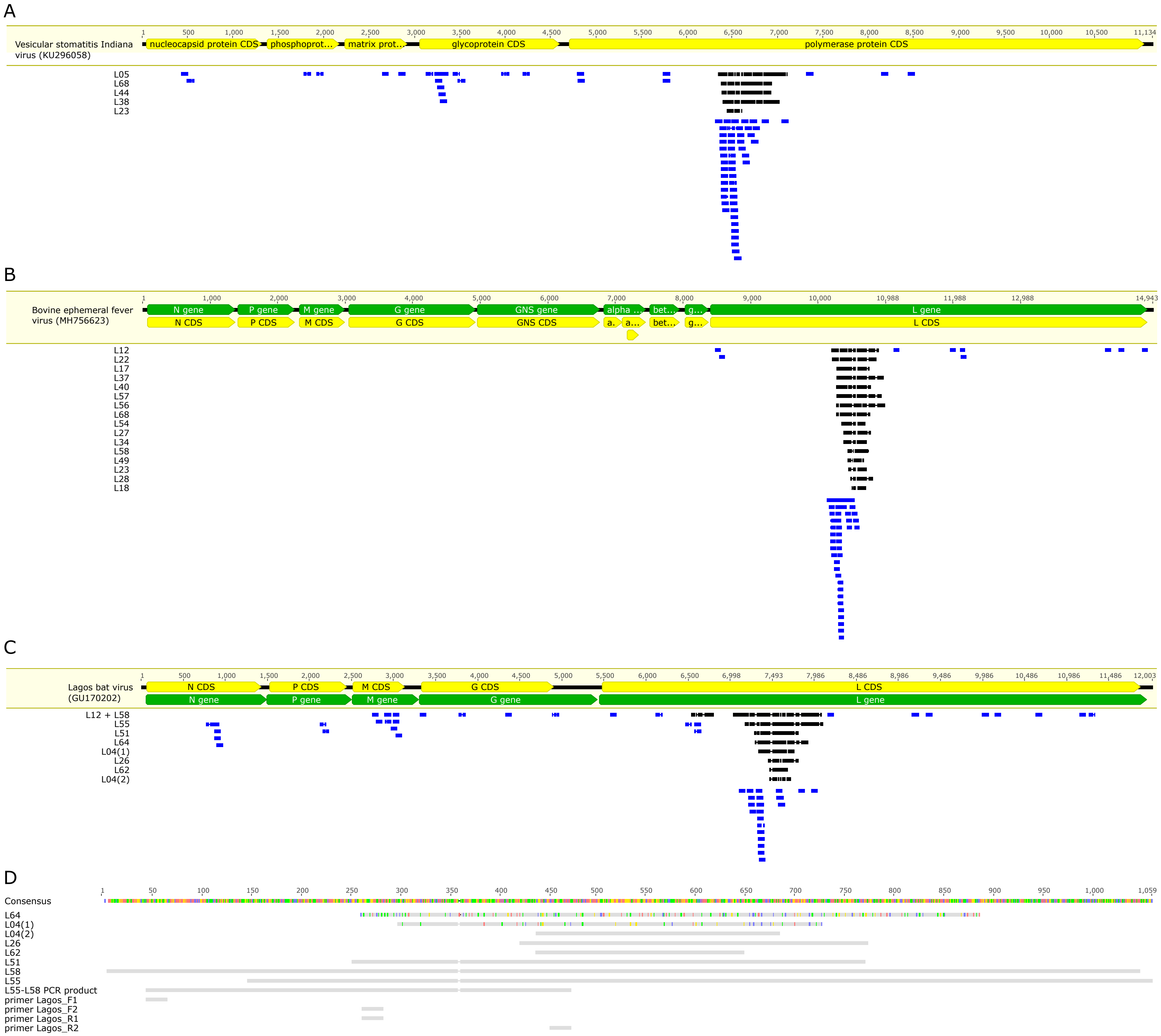
A
Vesicular stomatitis Indiana virus (KU296058)
nucleocapsid protein CDS
phosphoprot…
matrix prot…
glycoprotein CDS
polymerase protein CDS
L05
L68
L44
L38
L23
B
Bovine ephemeral fever virus (MH756623)
N gene
P gene
M gene
G gene
GNS gene
L gene
N CDS
P CDS
M CDS
G CDS
GNS CDS
L CDS
L12
L22
L17
L37
L40
L57
L56
L68
L54
L27
L34
L58
L49
L23
L28
L18
C
Lagos bat virus (GU170202)
N CDS
P CDS
M CDS
G CDS
L CDS
N gene
P gene
M gene
G gene
L gene
L12 + L58
L55
L51
L64
L04(1)
L26
L62
L04(2)
D
Consensus
L64
L04(1)
L04(2)
L26
L62
L51
L58
L55
L55-L58 PCR product
primer Lagos_F1
primer Lagos_F2
primer Lagos_R1
primer Lagos_R2

**Suppl. Fig. 4**

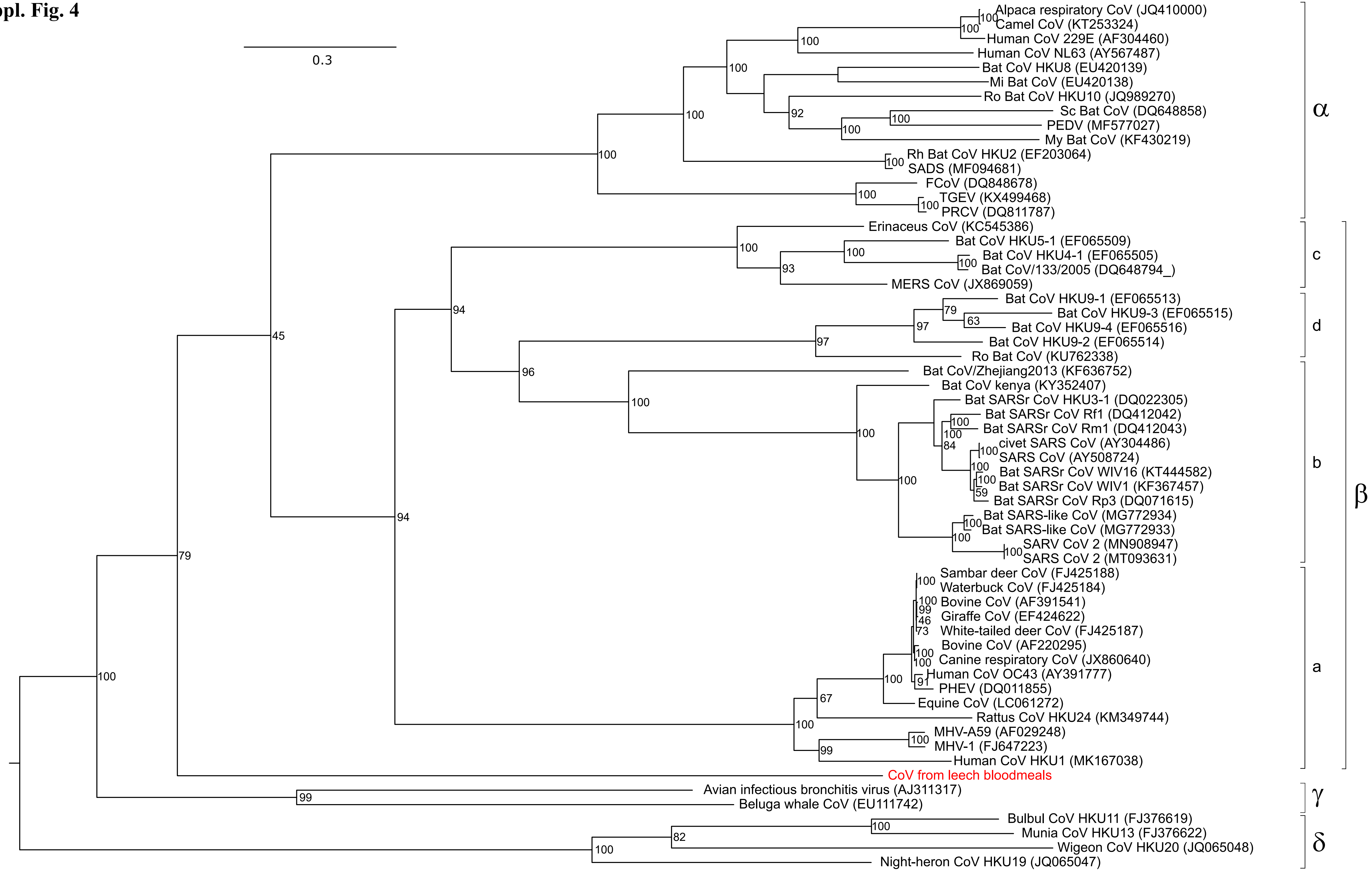
0.3
Alpaca respiratory CoV (JQ410000)
Camel CoV (KT253324)
Human CoV 229E (AF304460)
Human CoV NL63 (AY567487)
Bat CoV HKU8 (EU420139)
Mi Bat CoV (EU420138)
Ro Bat CoV HKU10 (JQ989270)
Sc Bat CoV (DQ648858)
PEDV (MF577027)
My Bat CoV (KF430219)
Rh Bat CoV HKU2 (EF203064)
SADS (MF094681)
FCoV (DQ848678)
TGEV (KX499468)
PRCV (DQ811787)
Erinaceus CoV (KC545386)
Bat CoV HKU5-1 (EF065509)
Bat CoV HKU4-1 (EF065505)
Bat CoV/133/2005 (DQ648794_)
MERS CoV (JX869059)
Bat CoV HKU9-1 (EF065513)
Bat CoV HKU9-3 (EF065515)
Bat CoV HKU9-4 (EF065516)
Bat CoV HKU9-2 (EF065514)
Ro Bat CoV (KU762338)
Bat CoV/Zhejiang2013 (KF636752)
Bat CoV kenya (KY352407)
Bat SARSr CoV HKU3-1 (DQ022305)
Bat SARSr CoV Rf1 (DQ412042)
Bat SARSr CoV Rm1 (DQ412043)
civet SARS CoV (AY304486)
SARS CoV (AY508724)
Bat SARSr CoV WIV16 (KT444582)
Bat SARSr CoV WIV1 (KF367457)
Bat SARSr CoV Rp3 (DQ071615)
Bat SARS-like CoV (MG772934)
Bat SARS-like CoV (MG772933)
SARV CoV 2 (MN908947)
SARS CoV 2 (MT093631)
Sambar deer CoV (FJ425188)
Waterbuck CoV (FJ425184)
Bovine CoV (AF391541)
Giraffe CoV (EF424622)
White-tailed deer CoV (FJ425187)
Bovine CoV (AF220295)
Canine respiratory CoV (JX860640)
Human CoV OC43 (AY391777)
PHEV (DQ011855)
Equine CoV (LC061272)
Rattus CoV HKU24 (KM349744)
MHV-A59 (AF029248)
MHV-1 (FJ647223)
Human CoV HKU1 (MK167038)
CoV from leech bloodmeals
Avian infectious bronchitis virus (AJ311317)
Beluga whale CoV (EU111742)
Bulbul CoV HKU11 (FJ376619)
Munia CoV HKU13 (FJ376622)
Wigeon CoV HKU20 (JQ065048)
Night-heron CoV HKU19 (JQ065047)
α
c
d
b
β
a
γ
δ

### Suppl. Fig. 5

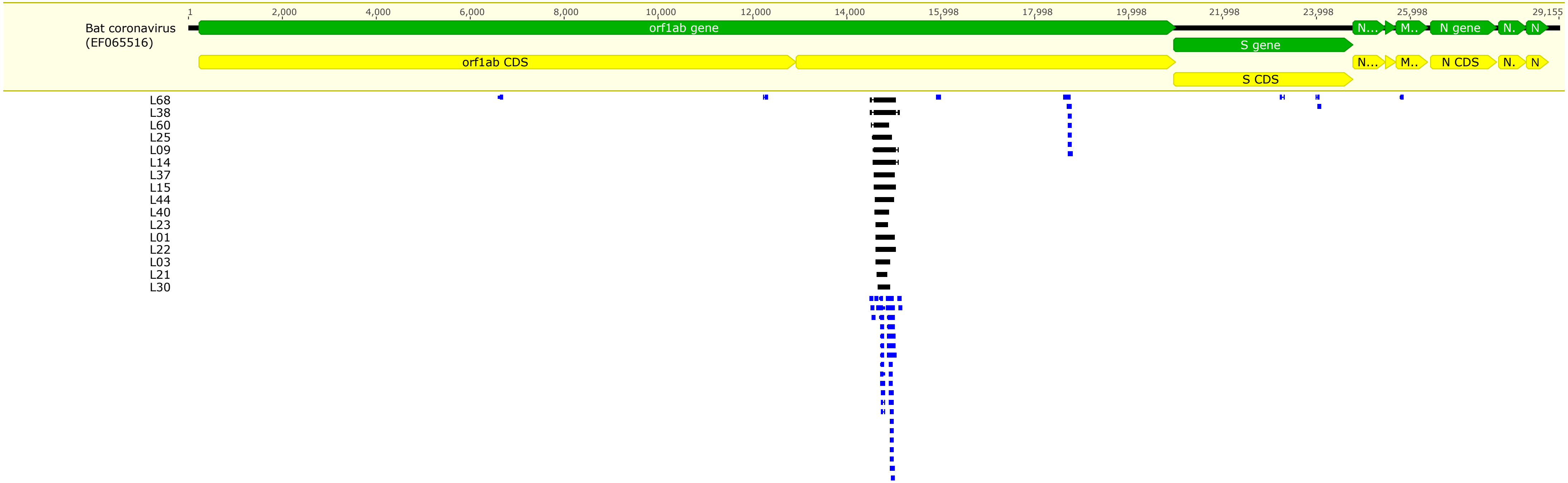

Bat coronavirus
(EF065516)
1
2,000
4,000
6,000
8,000
10,000
12,000
14,000
15,998
17,998
19,998
21,998
23,998
25,998
29,155
orf1ab gene
S gene
N gene
orf1ab CDS
S CDS
N CDS
L68
L38
L60
L25
L09
L14
L37
L15
L44
L40
L23
L01
L22
L03
L21
L30

### Suppl. Fig. 6

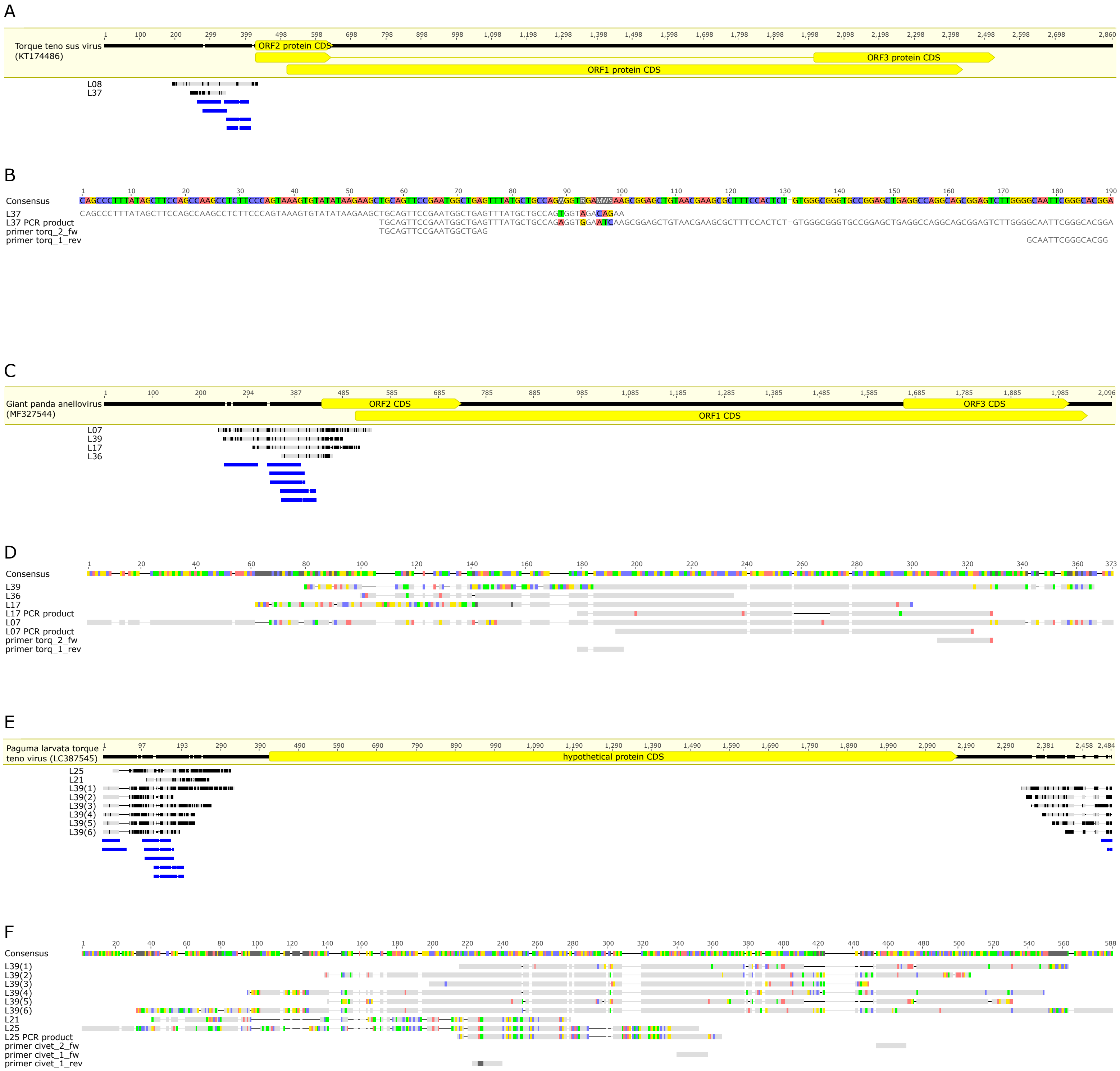
A
Torque teno sus virus (KT174486)
1 100 200 299 399 498 598 698 798 898 998 1,098 1,198 1,298 1,398 1,498 1,598 1,698 1,798 1,898 1,998 2,098 2,198 2,298 2,398 2,498 2,598 2,698 2,860
ORF2 protein CDS
ORF3 protein CDS
ORF1 protein CDS
L08
L37
B
1 10 20 30 40 50 60 70 80 90 100 110 120 130 140 150 160 170 180 190
Consensus CAGCCCTTTATAGCTTCCAGCCAAGCCTCTTCCCAGTAAAGTGTATATAAGAAGCTGCAGTTCCGAATGGCTGAGTTTATGCTGCCAGWGGTRGAMWSAAGCGGAGCTGTAACGAAGCGCTTTCCACTCT-GTGGGCGGGTGCCGGAGCTGAGGCCAGGCAGCGGAGTCTTGGGGCAATTCGGGCACGGA
L37 CAGCCCTTTATAGCTTCCAGCCAAGCCTCTTCCCAGTAAAGTGTATATAAGAAGCTGCAGTTCCGAATGGCTGAGTTTATGCTGCCAGTGGTAGACAGAA
L37 PCR product TGCAGTTCCGAATGGCTGAGTTTATGCTGCCAGAGGTGGAATCAAGCGGAGCTGTAACGAAGCGCTTTCCACTCT-GTGGGCGGGTGCCGGAGCTGAGGCCAGGCAGCGGAGTCTTGGGGCAATTCGGGCACGGA
primer torq_2_fw TGCAGTTCCGAATGGCTGAG
primer torq_1_rev GCAATTCGGGCACGG
C
Giant panda anellovirus (MF327544)
1 100 200 294 387 485 585 685 785 885 985 1,085 1,185 1,285 1,385 1,485 1,585 1,685 1,785 1,885 1,985 2,096
ORF2 CDS
ORF3 CDS
ORF1 CDS
L07
L39
L17
L36
D
1 20 40 60 80 100 120 140 160 180 200 220 240 260 280 300 320 340 360 373
Consensus
L39
L36
L17
L17 PCR product
L07
L07 PCR product
primer torq_2_fw
primer torq_1_rev
E
Paguma larvata torque teno virus (LC387545)
1 97 193 290 390 490 590 690 790 890 990 1,090 1,190 1,290 1,390 1,490 1,590 1,690 1,790 1,890 1,990 2,090 2,190 2,290 2,381 2,458 2,484
hypothetical protein CDS
L25
L21
L39(1)
L39(2)
L39(3)
L39(4)
L39(5)
L39(6)
F
1 20 40 60 80 100 120 140 160 180 200 220 240 260 280 300 320 340 360 380 400 420 440 460 480 500 520 540 560 588
Consensus
L39(1)
L39(2)
L39(3)
L39(4)
L39(5)
L39(6)
L21
L25
L25 PCR product
primer civet_2_fw
primer civet_1_fw
primer civet_1_rev

**Suppl. Fig. 7**

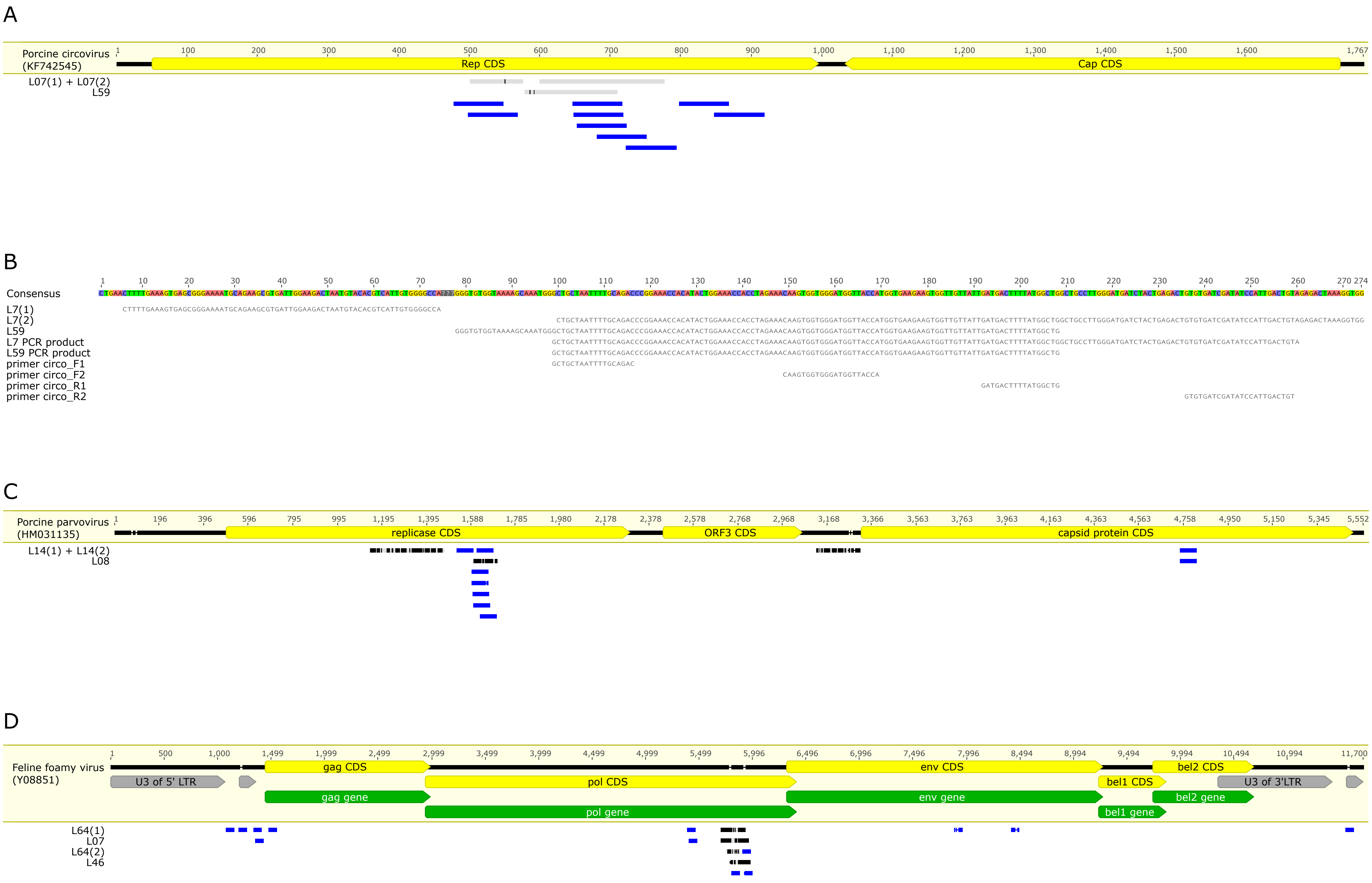
A
Porcine circovirus (KF742545)
Rep CDS
Cap CDS
L07(1) + L07(2)
L59
B
Consensus
L7(1)
L7(2)
L59
L7 PCR product
L59 PCR product
primer circo_F1
primer circo_F2
primer circo_R1
primer circo_R2
C
Porcine parvovirus (HM031135)
replicase CDS
ORF3 CDS
capsid protein CDS
L14(1) + L14(2)
L08
D
Feline foamy virus (Y08851)
U3 of 5' LTR
gag CDS
pol CDS
env CDS
bel1 CDS
bel2 CDS
U3 of 3'LTR
gag gene
pol gene
env gene
bel1 gene
bel2 gene
L64(1)
L07
L64(2)
L46

**Suppl. Fig. 8**

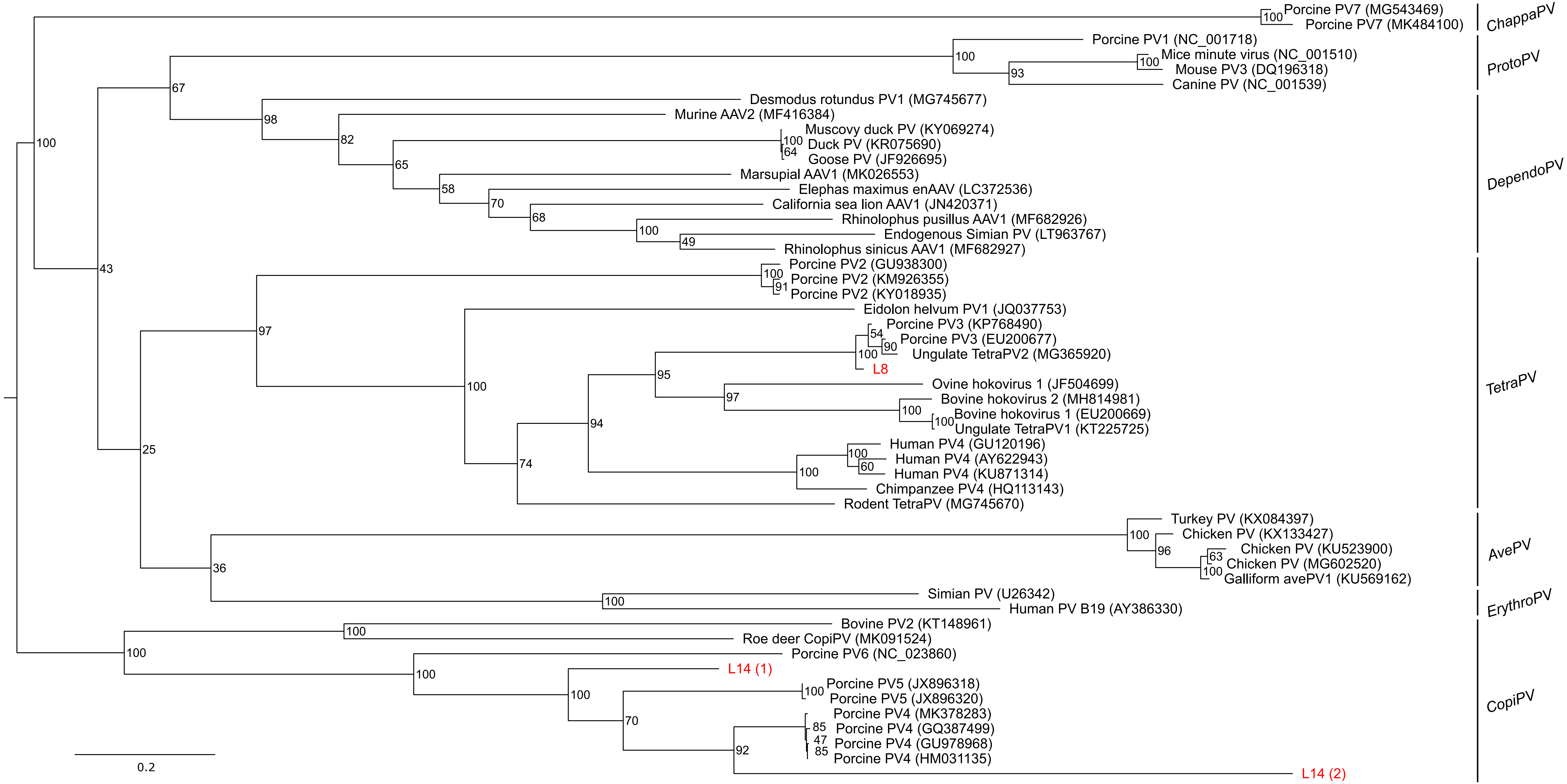

Porcine PV7 (MG543469)
Porcine PV7 (MK484100)
ChappaPV
Porcine PV1 (NC_001718)
Mice minute virus (NC_001510)
Mouse PV3 (DQ196318)
Canine PV (NC_001539)
ProtoPV
Desmodus rotundus PV1 (MG745677)
Murine AAV2 (MF416384)
Muscovy duck PV (KY069274)
Duck PV (KR075690)
Goose PV (JF926695)
Marsupial AAV1 (MK026553)
Elephas maximus enAAV (LC372536)
California sea lion AAV1 (JN420371)
Rhinolophus pusillus AAV1 (MF682926)
Endogenous Simian PV (LT963767)
Rhinolophus sinicus AAV1 (MF682927)
DependoPV
Porcine PV2 (GU938300)
Porcine PV2 (KM926355)
Porcine PV2 (KY018935)
Eidolon helvum PV1 (JQ037753)
Porcine PV3 (KP768490)
Porcine PV3 (EU200677)
Ungulate TetraPV2 (MG365920)
L8
Ovine hokovirus 1 (JF504699)
Bovine hokovirus 2 (MH814981)
Bovine hokovirus 1 (EU200669)
Ungulate TetraPV1 (KT225725)
Human PV4 (GU120196)
Human PV4 (AY622943)
Human PV4 (KU871314)
Chimpanzee PV4 (HQ113143)
Rodent TetraPV (MG745670)
TetraPV
Turkey PV (KX084397)
Chicken PV (KX133427)
Chicken PV (KU523900)
Chicken PV (MG602520)
Galliform avePV1 (KU569162)
AvePV
Simian PV (U26342)
Human PV B19 (AY386330)
ErythroPV
Bovine PV2 (KT148961)
Roe deer CopiPV (MK091524)
Porcine PV6 (NC_023860)
L14 (1)
Porcine PV5 (JX896318)
Porcine PV5 (JX896320)
Porcine PV4 (MK378283)
Porcine PV4 (GQ387499)
Porcine PV4 (GU978968)
Porcine PV4 (HM031135)
L14 (2)
CopiPV
0.2

Suppl. Fig. 9

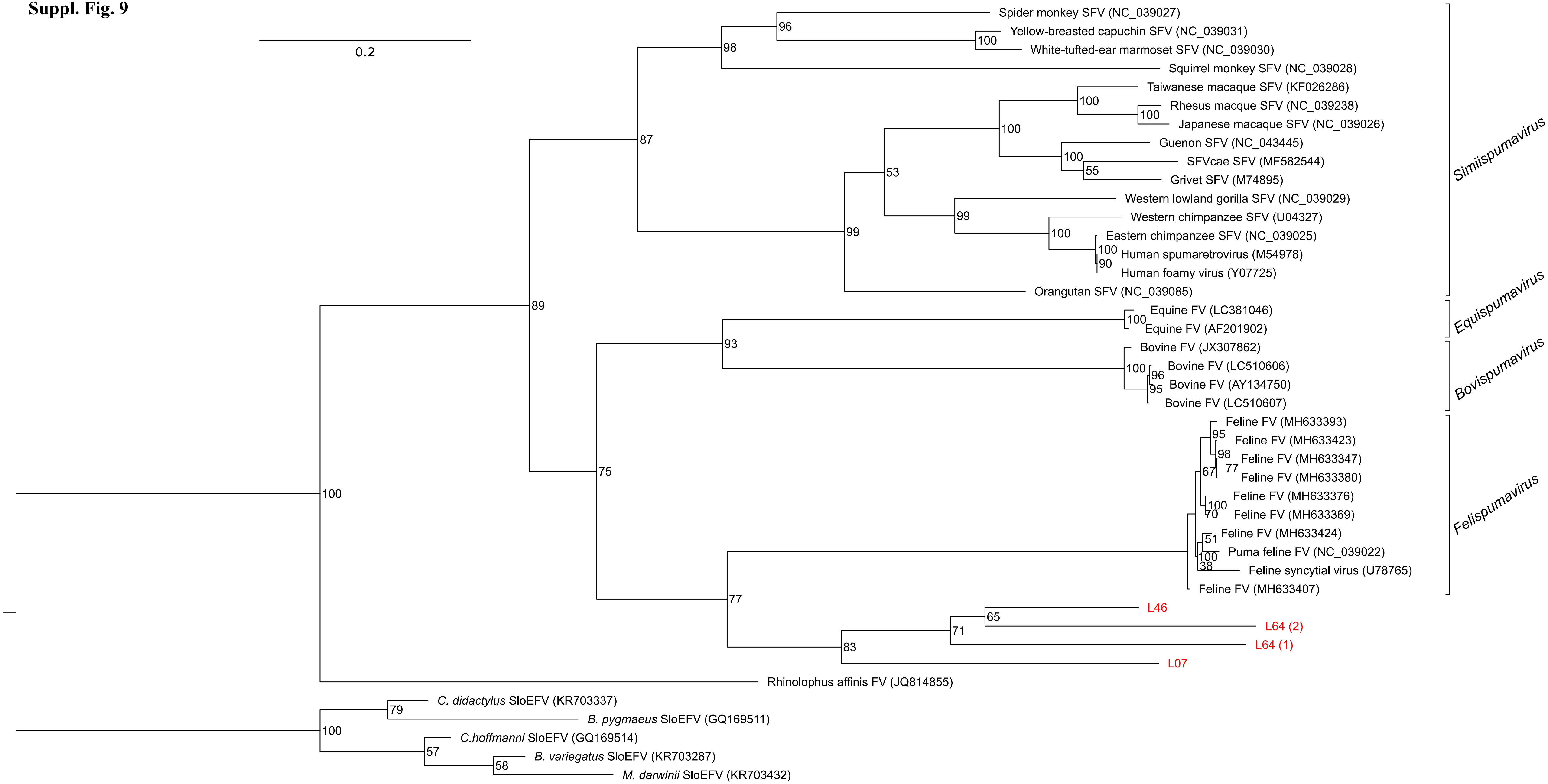
0.2
Spider monkey SFV (NC_039027)
Yellow-breasted capuchin SFV (NC_039031)
White-tufted-ear marmoset SFV (NC_039030)
Squirrel monkey SFV (NC_039028)
Taiwanese macaque SFV (KF026286)
Rhesus macque SFV (NC_039238)
Japanese macaque SFV (NC_039026)
Guenon SFV (NC_043445)
SFVcae SFV (MF582544)
Grivet SFV (M74895)
Western lowland gorilla SFV (NC_039029)
Western chimpanzee SFV (U04327)
Eastern chimpanzee SFV (NC_039025)
Human spumaretrovirus (M54978)
Human foamy virus (Y07725)
Orangutan SFV (NC_039085)
Equine FV (LC381046)
Equine FV (AF201902)
Bovine FV (JX307862)
Bovine FV (LC510606)
Bovine FV (AY134750)
Bovine FV (LC510607)
Feline FV (MH633393)
Feline FV (MH633423)
Feline FV (MH633347)
Feline FV (MH633380)
Feline FV (MH633376)
Feline FV (MH633369)
Feline FV (MH633424)
Puma feline FV (NC_039022)
Feline syncytial virus (U78765)
Feline FV (MH633407)
L46
L64 (2)
L64 (1)
L07
Rhinolophus affinis FV (JQ814855)
C. didactylus SloEFV (KR703337)
B. pygmaeus SloEFV (GQ169511)
C.hoffmanni SloEFV (GQ169514)
B. variegatus SloEFV (KR703287)
M. darwinii SloEFV (KR703432)
Simiispumavirus
Equispumavirus
Bovispumavirus
Felispumavirus

**Suppl. Fig. 10**

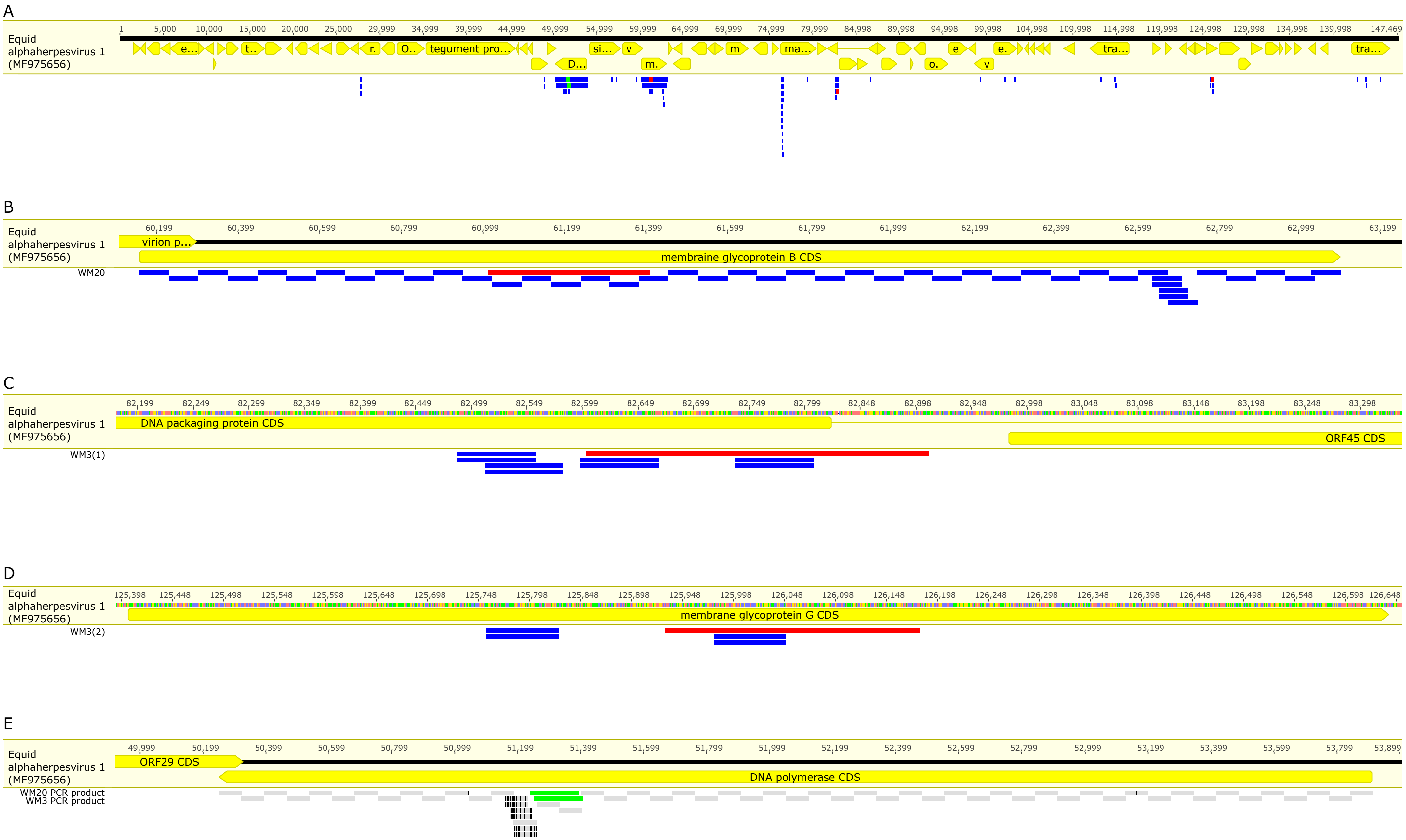
A
Equid alphaherpesvirus 1 (MF975656)
tegument pro...
B
Equid alphaherpesvirus 1 (MF975656)
virion p...
membraine glycoprotein B CDS
WM20
C
Equid alphaherpesvirus 1 (MF975656)
DNA packaging protein CDS
ORF45 CDS
WM3(1)
D
Equid alphaherpesvirus 1 (MF975656)
membrane glycoprotein G CDS
WM3(2)
E
Equid alphaherpesvirus 1 (MF975656)
ORF29 CDS
DNA polymerase CDS
WM20 PCR product
WM3 PCR product

### Suppl. Fig. 11

A

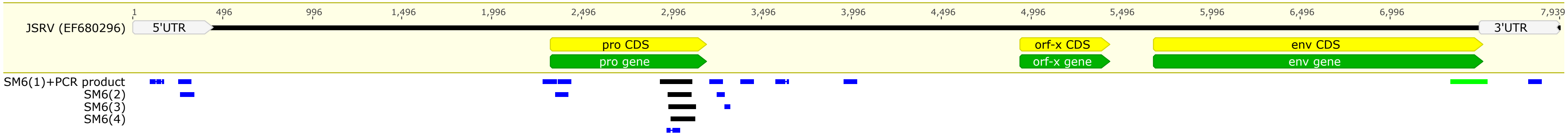


B

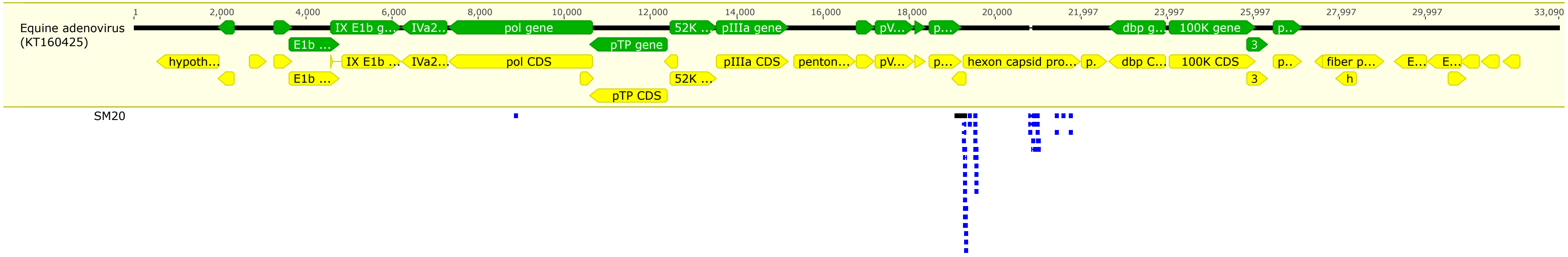


C

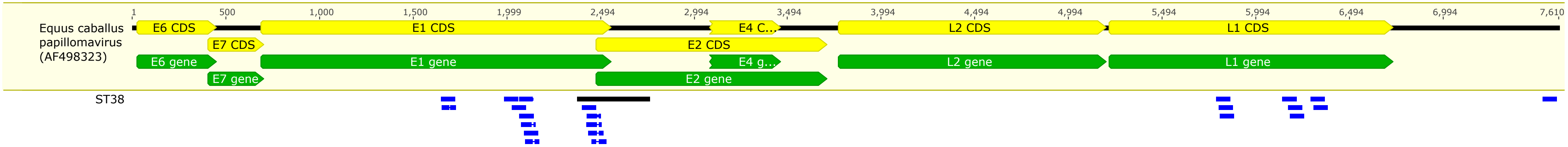

**Suppl. Fig. 12**

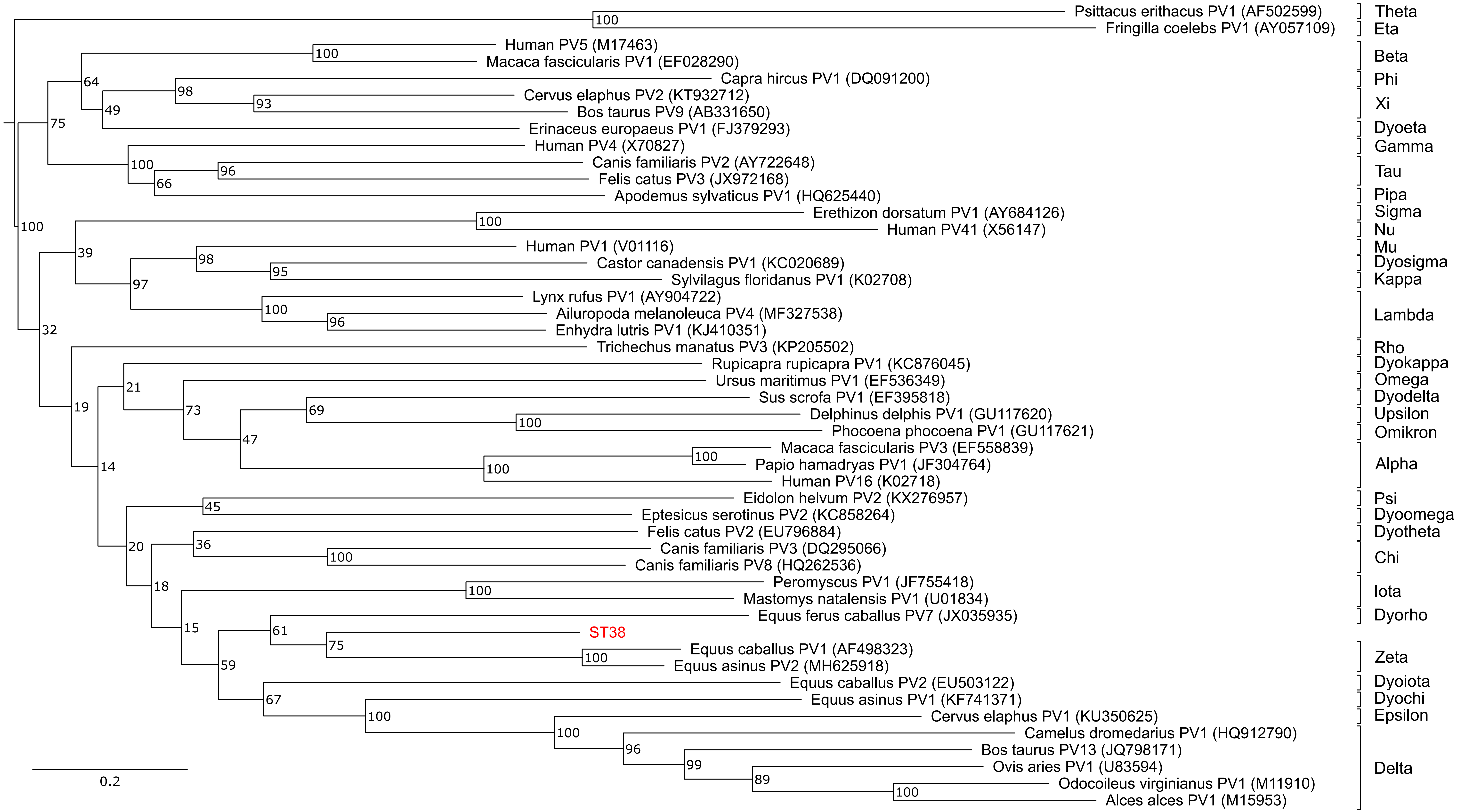
Psittacus erithacus PV1 (AF502599)
Fringilla coelebs PV1 (AY057109)
Human PV5 (M17463)
Macaca fascicularis PV1 (EF028290)
Capra hircus PV1 (DQ091200)
Cervus elaphus PV2 (KT932712)
Bos taurus PV9 (AB331650)
Erinaceus europaeus PV1 (FJ379293)
Human PV4 (X70827)
Canis familiaris PV2 (AY722648)
Felis catus PV3 (JX972168)
Apodemus sylvaticus PV1 (HQ625440)
Erethizon dorsatum PV1 (AY684126)
Human PV41 (X56147)
Human PV1 (V01116)
Castor canadensis PV1 (KC020689)
Sylvilagus floridanus PV1 (K02708)
Lynx rufus PV1 (AY904722)
Ailuropoda melanoleuca PV4 (MF327538)
Enhydra lutris PV1 (KJ410351)
Trichechus manatus PV3 (KP205502)
Rupicapra rupicapra PV1 (KC876045)
Ursus maritimus PV1 (EF536349)
Sus scrofa PV1 (EF395818)
Delphinus delphis PV1 (GU117620)
Phocoena phocoena PV1 (GU117621)
Macaca fascicularis PV3 (EF558839)
Papio hamadryas PV1 (JF304764)
Human PV16 (K02718)
Eidolon helvum PV2 (KX276957)
Eptesicus serotinus PV2 (KC858264)
Felis catus PV2 (EU796884)
Canis familiaris PV3 (DQ295066)
Canis familiaris PV8 (HQ262536)
Peromyscus PV1 (JF755418)
Mastomys natalensis PV1 (U01834)
Equus ferus caballus PV7 (JX035935)
ST38
Equus caballus PV1 (AF498323)
Equus asinus PV2 (MH625918)
Equus caballus PV2 (EU503122)
Equus asinus PV1 (KF741371)
Cervus elaphus PV1 (KU350625)
Camelus dromedarius PV1 (HQ912790)
Bos taurus PV13 (JQ798171)
Ovis aries PV1 (U83594)
Odocoileus virginianus PV1 (M11910)
Alces alces PV1 (M15953)
Theta
Eta
Beta
Phi
Xi
Dyoeta
Gamma
Tau
Pipa
Sigma
Nu
Mu
Dyosigma
Kappa
Lambda
Rho
Dyokappa
Omega
Dyodelta
Upsilon
Omikron
Alpha
Psi
Dyoomega
Dyotheta
Chi
Iota
Dyorho
Zeta
Dyoiota
Dyochi
Epsilon
Delta
0.2
